# scCross: A Deep Generative Model for Unifying Single-cell Multi-omics with Seamless Integration, Cross-modal Generation, and In-silico Exploration

**DOI:** 10.1101/2023.11.22.568376

**Authors:** Xiuhui Yang, Koren K. Mann, Hao Wu, Jun Ding

**Author notes:** Contributing authors.

## Abstract

Single-cell multi-omics illuminate intricate cellular states, yielding transformative insights into cellular dynamics and disease. Yet, while the potential of this technology is vast, the integration of its multifaceted data presents challenges. Some modalities have not reached the robustness or clarity of established scRNA-seq. Coupled with data scarcity for newer modalities and integration intricacies, these challenges limit our ability to maximize single-cell omics benefits. We introduce scCross: a tool adeptly engineered using variational autoencoder, generative adversarial network principles, and the Mutual Nearest Neighbors (MNN) technique for modality alignment. This synergy ensures seamless integration of varied single-cell multi-omics data. Beyond its foundational prowess in multi-omics data integration, scCross excels in single-cell cross-modal data generation, multi-omics data simulation, and profound in-silico cellular perturbations. Armed with these capabilities, scCross is set to transform the field of single-cell research, establishing itself in the nuanced integration, generation, and simulation of complex multi-omics data.

## Backgrounds

The advent of single-cell sequencing technologies has ushered in a new era of biological research, enabling scientists to deconvolve cellular heterogeneity in unprecedented detail [1–3]. This granularity has illuminated intricate cellular dynamics across myriad biological processes. To harness the full potential of this data deluge, a suite of computational tools has been developed, propelling advancements in fields as diverse as cancer biology, neurobiology, and drug discovery [4–8]. However, many of these tools are tailored to specific data modalities, such as scRNA-seq [9] or scATAC-seq [10, 11], often providing a piecemeal view of the cellular landscape.

As the single-cell sequencing paradigm matures, we are seeing the convergence of multiple data modalities, offering a holistic view of cellular states [3, 12–28]. Effective single-cell multi-omics data integration remains challenging, particularly when integrating unmatched data. Many single-cell multi-omics measurements are unmatched, profiling varied cell populations across modalities and thus missing the matched multimodal profiling of the same cells.

Existing methods have emerged to tackle this data integration challenge, each with its own set of advantages and drawbacks. Some, like Seurat [29], hinge on finding commonalities across modalities, while others explore nonlinear transformations or lean into deep learning. Many of these techniques face issues ranging from information loss [30] to noise susceptibility. Furthermore, due to the cost and difficulty in experiments, there is often a significant scarcity of matched single-cell multi-omics data that comprehensively profiles cellular states. Instead, we typically have abundant data for the most dominant modality (like scRNA-seq) but limited or no profiling for other modalities (e.g., epigenetics). The presence of other matched single-cell multi-omics data offers the potential for cross-modal generation of missing modalities from the more abundant ones by transferring knowledge learned from the reference.

Here, we introduce scCross. At its foundation, scCross excels in its function of single-cell multi-omics data integration, bringing unparalleled precision to the assimilation of diverse data modalities. While preserving its primary proficiency in this arena, the most distinctive feature of scCross emerges: its prowess in cross-modal single-cell data generation. This capability, layered atop its integration strength, unlocks transformative potentials for researchers to bridge disparities between abundant and scant modalities. Building on these two principal functionalities, scCross further showcases its versatility by simulating single-cell multi-omics data with high fidelity. Moreover, the tool unveils opportunities for in-silico perturbations within and across modalities, enabling researchers to postulate and evaluate potential cellular interventions. By capturing and elucidating intricate cellular states, scCross equips the scientific community with a thorough grasp of cellular dynamics across modalities. scCross stakes its claim as a vanguard in single-cell multi-omics integration, generation, and simulation.

## Results

### Overview of scCross

scCross employs the Variational utoencoder(VAE)-Generative adversial network(GAN) deep generative framework to integrate single-cell multi-omics datasets, generate cross-modal data, simulate multi-omics data, and perform in-silico perturbations within and across modalities(Fig. 1 and Supplementary Fig. S1). The methodology begins by training modality-specific Variational Autoencoders (VAEs) to capture low-dimensional cell embeddings for each data type, supplemented by the integration of gene set score vectors as representative features. With these embeddings learned, they are integrated into a common latent space. The harmonization of the modalities’ data distributions then ensues. The Generative Adversarial Network (GAN) refines this process, with a discriminator network juxtaposed against the VAE generator. This structure ensures a robust multi-modal data integration and confirms the alignment between the actual and generated single-cell inputs.

**Fig. 1.**
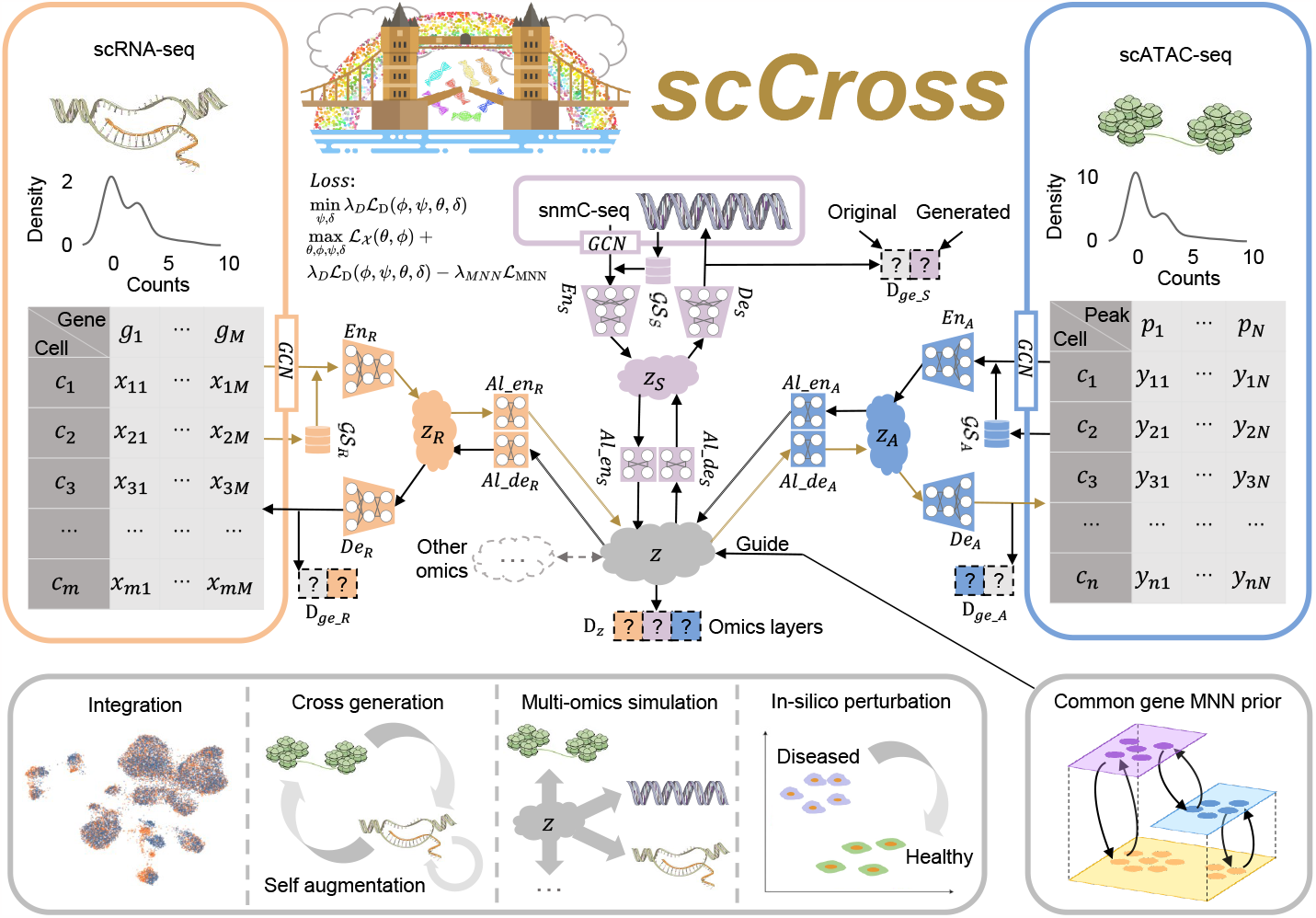
Architecture of the scCross framework. scCross employs modality-specific Variational Autoencoders to capture cell latent embeddings **z**_*R*_, **z**_*S*_, **z**_*A*_, …, for each omics type. During single-cell data integration, the method leverages biological priors by integrating gene set matrices 𝒢 𝒮_*R*_, 𝒢 𝒮_*S*_, 𝒢 𝒮_*A*_, …, as additional features for each cell. The framework then harmonizes these enriched embeddings into shared embeddings *z* using further variational autoencoders and critically, bidirectional aligners. Bidirectional aligners are pivotal for the cross-modal generation, depicted by brown arrows signifying the transition from scRNA-seq to scATAC-seq. Mutual Nearest Neighbor (MNN) cell pairs ensure precision during alignment. Discriminator D_*z*_ maintains the integration of different omics and discriminators 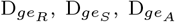, …, maintain the integrity and consistency of generated data. scCross offers a powerful toolbox for single-cell data integration, facilitating cross-modal data generation, single-cell data self augmentation, single-cell multi-omics simulation, and in-silico perturbations, making it versatile for a plethora of single-cell multi-omics challenges.

Venturing beyond mere integration and other functions, the model is specifically designed to facilitate cross-modal data generation. The bidirectional aligner is crucial for this cross-modal generation, as it decodes the shared latent embedding into a distinct modality, relying on the cell embedding sourced from the other modalities. This ambition of the model requires a dual alignment: a congruent data distribution and a meticulous coordination of individual cells across modalities. Mutual Nearest Neighbor (MNN) cell pairs are strategically employed as alignment anchors, guiding this intricate process. The optimization of the neural network aims to minimize discrepancies across modalities, especially those evident between MNN pairs. Once trained, the model enables the generation of single-cell data across modalities, encoding data from one modality into the latent space, and subsequently decoding it into another. It can also simulate multi-omics data and facilitates in-silico perturbations within and across modalities, revealing potential modulations of cellular states. By merging single-cell multi-omic data into a unified latent space and enabling information flow across modalities, scCross sets the stage for various multi-omic tasks.

### scCross empowers cross-modal single-cell data generation

Single-cell multi-omics data offers deep insights into the complexities of cellular states. Yet, factors such as cost and experimental challenges often impede our capability to fully harness its advantages, leading to some modalities being less explored than others. In this context, scCross emerges as a crucial tool, focused on bolstering cross-modal single-cell data generation to tap into the vast potential of single-cell multi-omics. It achieves this by seamlessly integrating knowledge across modalities, drawing from established reference single-cell multi-omics datasets encompassing all desired modalities or from multi-omic datasets where the underrepresented modalities were insufficiently profiled.

While tools like scglue have their merits in single-cell multi-omics data integration, they inherently lack the capability for cross-modal generation. Specifically, scglue’s entire framework emphasizes its unsuitability for crossmodal data generation, especially when noting that its decoder is designed to process embeddings from the input of the same modality. Contrastingly, scCross is architected to bridge such analytical gaps. To showcase its prowess and endorse its robustness and evaluate the method’s performance in situations with limited or partial single-cell data of another modality of interest, we divided the matched mouse cortex dataset [31] into two parts — 20% for training and the remainder for testing. Leveraging the model trained on the limited matched (20%) single-cell data, we applied the method to cross-generate the single-cell RNA-seq for the other missing 80% single-cell RNA-seq. The results, depicted in Fig. 2a-e, shed light on scCross’s ability not only to generate but also to uphold the biological nuances of the original data. The UMAP visualization [32] in Fig. 2a showcases both the original and generated scRNA-seq data clustering closely, hinting at a high similarity in their distributions. A compelling confirmation of this is observed in Fig. 2b, where the cell type proportion between the original and generated datasets demonstrates a correlation of 0.89 with a *p*-value of 0.001311. In Fig. 2c, the gene expression of the top 5 genes for each cell type between both datasets remains largely consistent, with a correlation of 0.93 and a *p*-value of 5.356 *×* 10^*−*177^. The remarkable similarity between the actual data and the crossed modality-generated data is apparent in those downstream analysis outcomes. Cell-cell interaction patterns, when analyzed through the CellChat tool [33] as seen in Fig. 2d, not only closely emulate those in the original dataset but also provide a window into potential interactions, leveraging insights from the scATAC-seq data of the other modality. This is quantitatively expressed by a correlation coefficient of 0.82 and a compellingly significant *p*-value of 1.356 *×* 10^*−*20^. Furthermore, pathway analysis based on the top 100 marker genes for each cell type, conducted with the ToppGene platform [34] and depicted in Fig. 2e, reinforces the crossed data’s relevance for robust biological interpretation. This analysis maintains a high correlation of 0.85 with a *p*-value of 2.531 *×* 10^*−*13^. The integrity of the crossed data is further substantiated by UMAP analyses presented in Supplementary Figs. S2-S5, which cover a comprehensive set of benchmark datasets, both matched and unmatched. Collectively, these analyses affirm the crossed data’s exceptional ability to mirror the biological intricacies and preserve the analytical properties of the actual data.

**Fig. 2.**
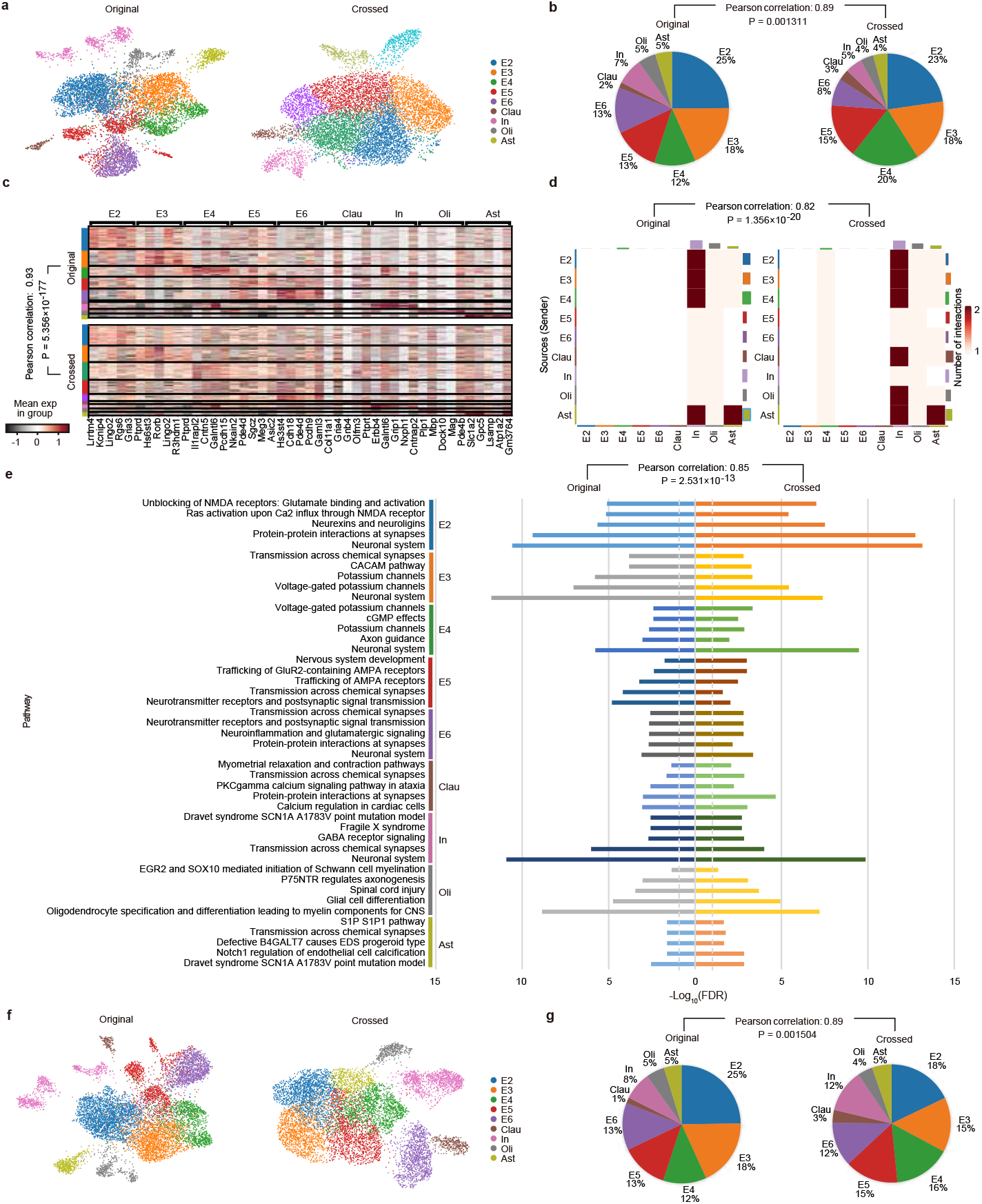
Performance of scCross in crossmodal single-cell data generation: emphasis on similarity between actual and cross-generated data. **a-e**, Results when the model was trained on the partial dataset: **a**, UMAP visualization emphasizes the overlapping distribution between the original and cross-generated scRNA-seq data. **b**, Comparison of cell type proportions reveals a robust correlation of 0.89 (*P*=0.001311). **c**, Expression levels of the top 5 genes per cell type exhibit a correlation of 0.93 (*P* = 5.356 *×* 10^*−*177^). **d**, Cell-cell interaction metrics show congruity across cell types with a correlation of 0.82 (*P* = 1.356 *×* 10^*−*20^). **e**, Ratios of genes in the most significant pathways for each cell type validate a correlation of 0.85 (*P* = 2.531 *×* 10^*−*13^) between the original and cross-generated datasets. **f-g**, Results when the model was trained on independent reference multi-omics datasets: **f**, UMAP visualization illustrates the clustering resemblance between the original and cross-generated matched mouse scRNA-seq data. **g**, Cell type proportions underscore a correlation of 0.89 (*P*=0.001504) between the original and cross-generated scRNA-seq data.

scCross’s capability for cross-modality generation was further rigorously tested in a scenario devoid of any direct training set for the missing modality. Utilizing an unmatched reference single-cell multi-omics dataset containing both RNA-seq and ATAC-seq data from the same samples [35], scCross was trained to harness and transfer this integrated knowledge. It was then tasked with generating single-cell RNA-seq data from single-cell ATAC-seq data alone, sourced from another matched multi-omics dataset [31]. The authenticity of this cross-generated RNA-seq data was critically assessed against the actual RNA-seq dataset, with the UMAP overlap in Fig. 2f serving as a visual testament to scCross’s crossmodal generation precision. The correlation in cell type composition between the cross-generated RNA-seq data and the actual RNA-seq data further substantiates scCross’s performance, achieving a correlation of 0.89 and a *p*-value of 0.001504 as illustrated in Fig. 2g. The practicality and relevance of scCross’s output are reinforced through various downstream analyses, detailed in Supplementary Fig. S6. This showcases scCross’s ability to not just mimic, but also potentially extrapolate and fill in gaps in single-cell omics profiles, reinforcing its role as a potent tool for advancing biological insights in the absence of comprehensive multi-omic datasets.

Beyond the cross-generational capabilities from one modality to another, our exploration extended to augment the inputs within the same modality, leveraging the inherent reconstruction and augmentation strength of the VAE-GAN framework [36–38]. This captivating line of inquiry uncovered that the data underwent an augmentation phase, which in turn enriched its quality and expanded its informational depth. Detailed results and visualizations can be found in Supplementary Fig. S7.

### Simulation of matched single-cell multi-omics data

Another capability of scCross is its ability to generate matched single-cell multi-omics data. This computational prowess opens the door to numerous practical applications. One immediate use is in benchmarking existing single-cell multi-omics integration methods, especially in scenarios where true matched single-cell multi-omics data is lacking for accurate evaluations. Moreover, the simulated data can also serve as a basis for deciphering intricate inter-omics relationships, predicting states of unmeasured omics, or filling in the gaps in studies with incomplete modalities.

One of the standout features of scCross is its tailored generation capability, targeting specific cell types of interest across various omics layers. This becomes indispensable when focusing on rare cell populations, which often are underrepresented in sequenced datasets. The limited number of cells in these populations poses challenges for in-depth analysis and examination. Here, scCross provides a valuable solution. By simulating data up to 5X the original count, as shown in Panels 3a and 3b, scCross can substantially upscale the representation of these rare populations, enabling more robust and comprehensive statistical analyses.

Panel 3a illustrates the simulated single-cell RNA-seq data of Ast cells in matched mouse cortex dataset, both at its original count and a 5X upscale with the whole original matched mouse cortex dataset as cluster background. Panel b mirrors this information for the scATAC-seq data. The high-quality cell clustering shown in these panels testifies to the effectiveness of our simulation approach. Other cell types in matched mouse cortex dataset also give out the same results (Supplementary Fig. S8).

To showcase the efficacy of scCross in simulating single-cell multi-omics data for rare cell populations, we targeted the Astrocyte (Ast) cells within the dataset, augmenting their representation by a fivefold increase. This upscaling was performed to better understand the potential implications of such rare cell types in subsequent analyses. We then conducted a comparative study between the original and the simulated dataset, centering on the enriched Gene Ontology (GO) terms and pathways linked to the Ast cell marker genes prior to and following their expansion. In Fig. 3c, the enriched terms are ranked and displayed, with particular attention to the leading terms. The two principal terms, ‘Vascular transport’ [39] and ‘Transport across blood-brain barrier’ [40], were found to be closely associated with the physiological roles of the Ast cells. The pathway ‘NOTCH1 regulation of endothelial cell calcification’ shows significance only in the 5x upscaled Ast cell simulated data, but not in the original dataset. These findings were verified to be associated with the signaling processes in Ast cells [41], thereby further substantiating the potential effectiveness of scCross in enhancing rare cell populations. Further validating the simulation’s accuracy, Fig. 3d compares the simulated data against the actual dataset for these rare Ast cells. The RRHO plot therein confirms that scCross preserves the key attributes of the Ast cells, thereby substantiating the simulated data’s fidelity for this specific subset within the larger dataset.

**Fig. 3.**
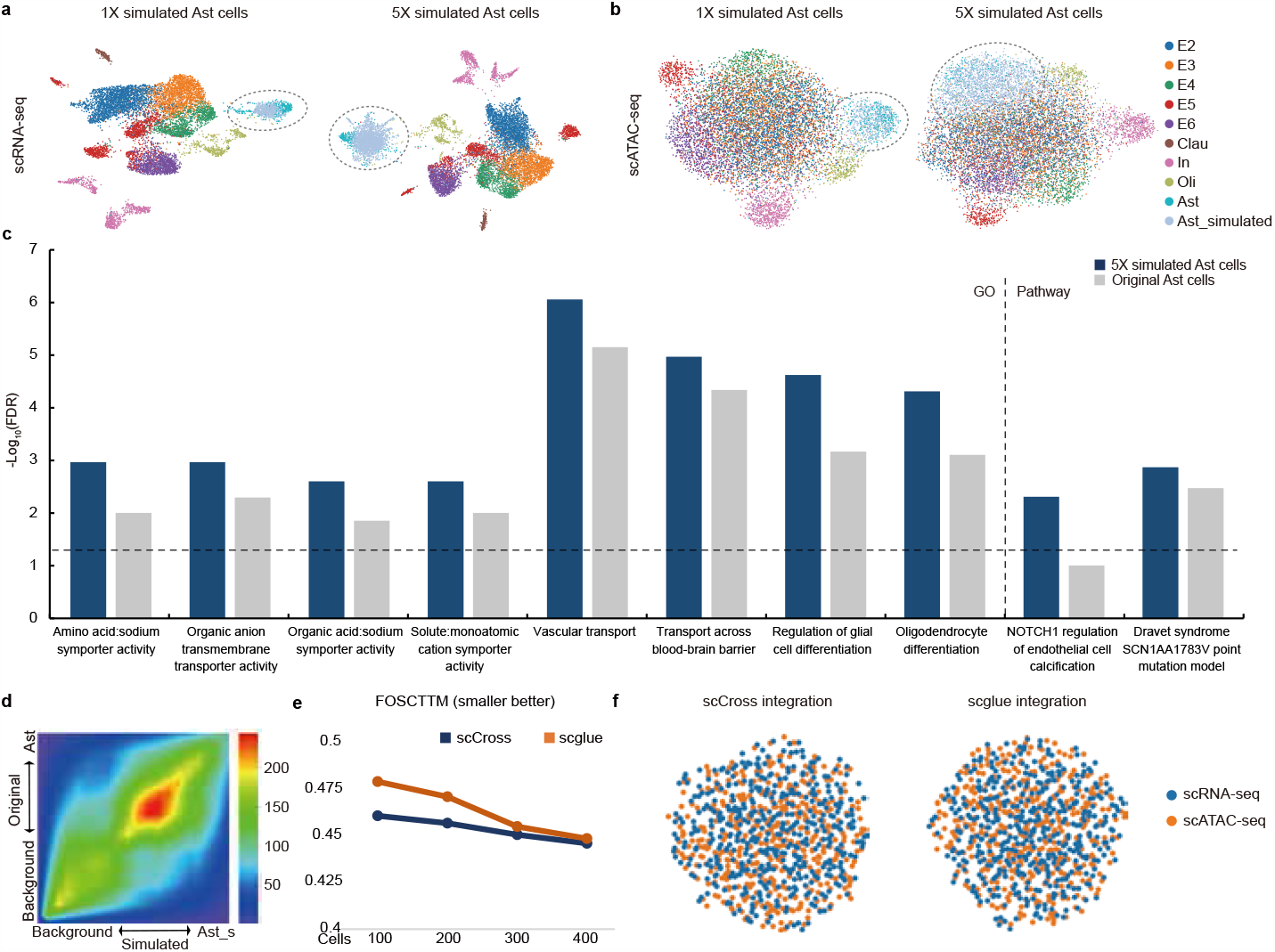
Simulation of matched single-cell multi-omics data. **a**, UMAP visualization showcasing the prowess of scCross in simulating the scRNA-seq data of rare Ast cells in matched mouse cortex dataset (highlighted by gray cycles), presented at 1X and 5X their original count with the whole original dataset as cluster background. The 5X scale highlights the model’s capability to upscale the representation of underrepresented populations like Ast cells. **b**, Parallel to Panel a, this UMAP visualization delineates the simulated scATAC-seq data for Ast cells at the same 1X and 5X scales (highlighted by gray cycles), reinforcing the model’s consistent performance across omic modalities. **c**, The analysis contrasts the Gene Ontology (GO) terms and pathways enriched among the top biomarkers of the original Astrocyte (Ast) cells (scRNA-seq) with those of its 5X augmented simulated equivalent. The comparison underscores the model’s proficiency in amplifying key biological signals within the framework of multi-omics data simulation. **d**, RRHO plot emphasizing the significant correlation between the original Ast scRNA-seq data and its 1X simulated counterpart. This highlights the model’s accurate capture of key biomarker genes during the simulation. **e**, FOSCTTM integration metrics underscore the potential of the 1X simulated Ast cells’ single-cell multi-omics data (encompassing both scRNA-seq and scATAC-seq) as an evaluative benchmark. Both scCross and scglue are compared; however, the inherent bias towards scCross due to the data’s origin should be acknowledged. **f**, UMAP visualization displaying the robust cell mixing achieved by both scglue and scCross when processing the 1X simulated Ast cells’ single-cell multi-omics data, providing a vivid demonstration of their respective capabilities.

The utility of this simulated data extends beyond mere generation. Panels 3e and 3f spotlight its potential applications. The panels underscore that the generated single-cell multi-omics data (1X simulated Ast) are “matched” since cells from distinct modalities derive from a consistent joint latent space. Panel e showcases the potential of using simulated data as an effective ground-truth to appraise single-cell data integration techniques. This perspective is reinforced in Panel f’s UMAP scatter plot, revealing exceptional cell mixing across modalities by both scglue and scCross. However, when examining the comparison in Panel e, caution is advised. Since these simulated datasets originate from the scCross model, there might be a slight bias in favor of scCross. Nevertheless, this comparison underscores that these simulated datasets can serve as robust matched single-cell multi-omics benchmarks for examining a range of other integration methods especially when ground truth is experimentally unattainable.

### scCross enables multi-omic in-silico perturbation for exploring potential cellular state intervention

In our endeavor to showcase the capabilities of scCross, we highlight its expertise in conducting in-silico multi-omic perturbations. These capabilities not only enable the exploration of potential intervention strategies for cellular state manipulation but also provide insights into the intricacies of molecular responses, particularly those seen in COVID-19 infections using the single-cell multi-omic dataset derived from a comprehensive study [17].

We began by measuring the scaled cosine distance between the in-silico perturbed latent matrix of COVID-19 cells and that of healthy cells (Fig. 4a) in the joint latent space produced by scCross. This allowed us to pinpoint signature marker genes that bridge the these two conditions (i.e., diseased vs. healthy) via in-silico perturbations (see Methods for details). The identified critical disease-associated marker genes from this in-silico perturbation, presented in Fig. 4b, are in line with recent findings. A notable observation was the down-perturbation of CCL3 in COVID-19 samples, suggesting its overexpression in COVID-19 patients. This concurs with prior studies that identified CCL3’s role with inflammatory macrophages and its contribution to inflammatory tissue damage and respiratory issues in COVID-19 patients [42–44]. Furthermore, the GO and pathway outcomes from the down-perturbation gene findings, showcased in Fig. 4c, unveiled significant associations with immune responses. Aspects like signaling receptor binding, immunoglobulin receptor binding, antigen binding, humoral immune response, adaptive immune response, and immunoglobulin complex are highlighted. These observations align well with the recognized impact of immune responses and chemokine activations in the context of COVID-19 [43]. The performance on the COVID-19 adverse outcome pathway [45], Network map of SARS-CoV-2 signaling pathway [46], and SARS-CoV-2 innate immunity evasion and cell-specific immune response pathway [47] further validates scCross’s potential in pinpointing signature genes between disease conditions within the same modality.

**Fig. 4.**
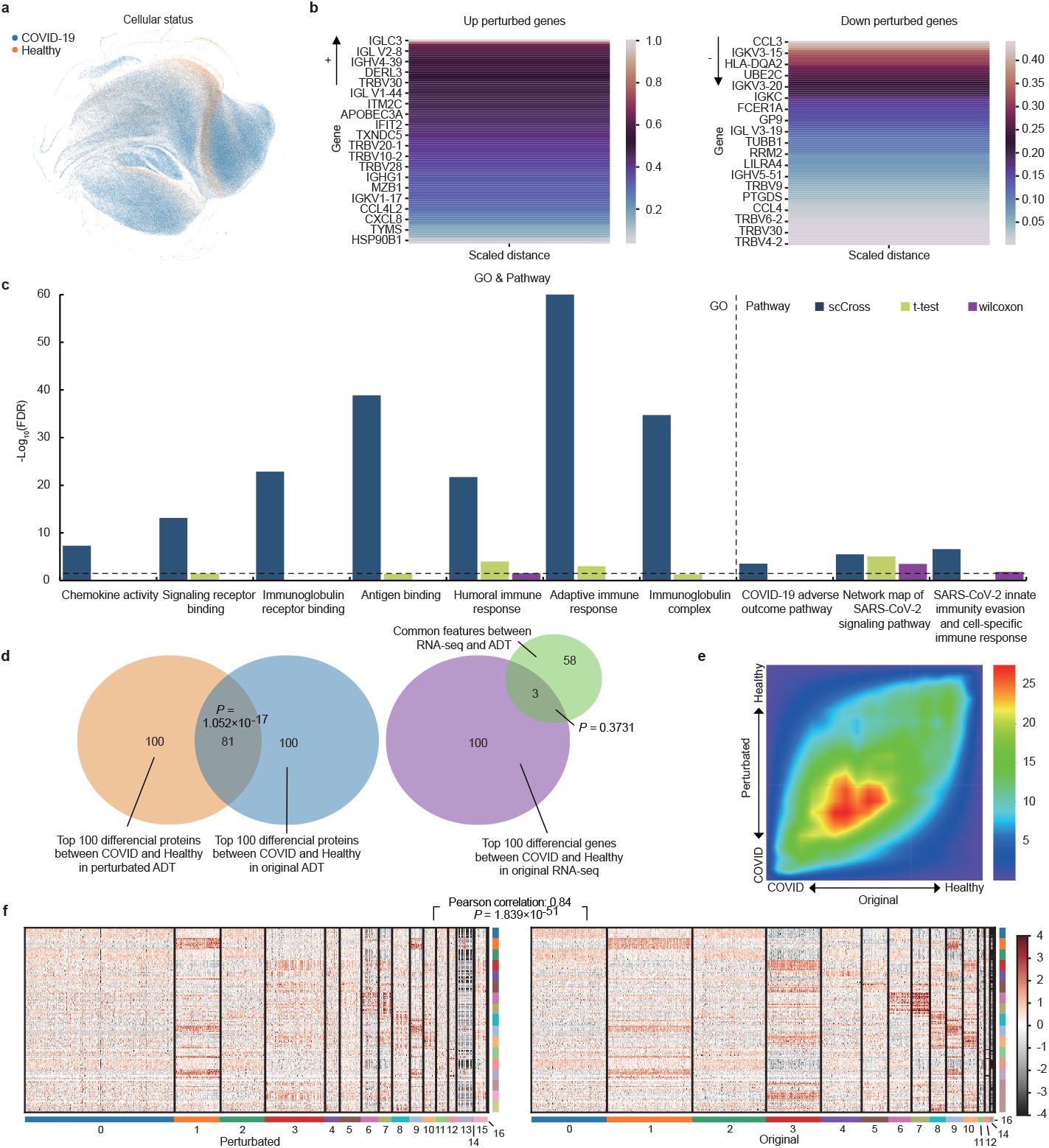
scCross empowers in-silico multi-omic perturbation as demonstrated in the COVID-19 dataset. **a**, UMAP visualization of COVID-19 scRNA-seq and ADT data clusters. **b**, Signature genes for in-silico perturbation bridging COVID-19 and healthy states. **c**, Pathways and GO terms of the top differential genes detected by scCross (down-perturbation gene findings), with -log(*p*-value) indicating significance, compared with t-test and wilconxon. **d**, Gene comparison between scCross-derived perturbed protein data and common-gene based on scRNA-seq to protein translation. **e**, RRHO plot comparing cross-modal perturbed and original COVID-19 protein data against healthy samples. **f**, Heatmap aligning original and scCross-derived perturbed COVID-19 protein data.

Fig. 4d delves into scCross’s potential in performing cross-modality insilico perturbations. By specifically perturbing the gene expression of signature genes that potentially transition healthy cells to a COVID-19-like state in the healthy scRNA-seq data and utilizing cross-generation, we generate perturbed protein data (ADT) across modalities to simulate the COVID condition. This perturbed ADT, when juxtaposed with the healthy protein data, yielded differential proteins (List A). This list was then contrasted with the actual differential protein list (List B) obtained from single-cell ADT comparisons between healthy and COVID datasets. The profound overlap between the two lists showcases the accuracy of cross-modal in-silico perturbations using scCross. Only three of the top 100 differentially expressed genes in scRNA-seq corresponded with those identified in single-cell protein measurements. This misalignment underscores the pitfalls of the naive strategy, which assumes that a gene differential at the RNA level is similarly differential at the protein level. Such discrepancies emphasize the efficacy and critical importance of scCross in these analyses.

Fig. 4e, with its RRHO plot, further magnifies the similarity between the cross-modal perturbed ADT and the actual ADT COVID dataset. The heatmap in Fig. 4f underlines this alignment, with a *p*-value of 1.839 *×* 10^*−*51^ (via the Wilcoxon test) cementing the proximity between the perturbed protein data and actual single-cell protein measurements. Conclusively, the data presents a compelling case for scCross’s ability to manage in-silico perturbations across modalities, reiterating its value in intricate multi-omic cellular state explorations.

### scCross improves single-cell multi-omics integration over established methods

Single-cell multi-omics data has emerged as a transformative tool that comprehensively captures cellular states, offering profound insights into cellular identity and function. Effective integration of the multi-omics data produces cell embeddings crucial for diverse analyses. These embeddings are pivotal for precise cell clustering and identification, and they support numerous downstream tasks. Metrics such as the Adjusted Rand Index (ARI) [48] and Normalized Mutual Information (NMI) [49] are invaluable in evaluating the quality of these embeddings, reflecting the proficiency of an integration method in retaining biological information.

We benchmarked scCross against several state-of-the-art single-cell data integration methods, including Seurat v4 [29], scglue [36], and uniPort [37]. Our analysis utilized three gold-standard datasets derived from recent single-cell multi-omics sequencing, encompassing simultaneous scRNA-seq and scATAC-seq profiling. These datasets are matched mouse cortex [31], matched mouse lymph nodes [50], and matched mouse atherosclerotic plaque immune cells (GSE240753). Additionally, we benchmarked the methods on another unmatched dataset, which includes scRNA-seq [35], scATAC-seq sourced from the 10X Genomics website, and an snmC-seq dataset [18].

Against this landscape, scCross stands as a leading method in single-cell multi-omics data integration, showcasing superior performance relative to other methods (Fig. 5 and Supplementary Figs. S9-S12). This distinction is evident from two primary angles. Firstly, scCross excels in downstream cell clustering, as demonstrated by its outstanding ARI, NMI, and cell type average silhouette width [36, 51] metrics across all benchmark datasets (Fig. 5a). Secondly, its prowess in cell mixing is apparent, seamlessly blending cells from different modalities. Effective cell mixing is indicative of high-quality integration and the method’s ability to represent the intricate nuances of each modality. Please refer to the Methods for a detailed description of the evaluation metrics.

**Fig. 5.**
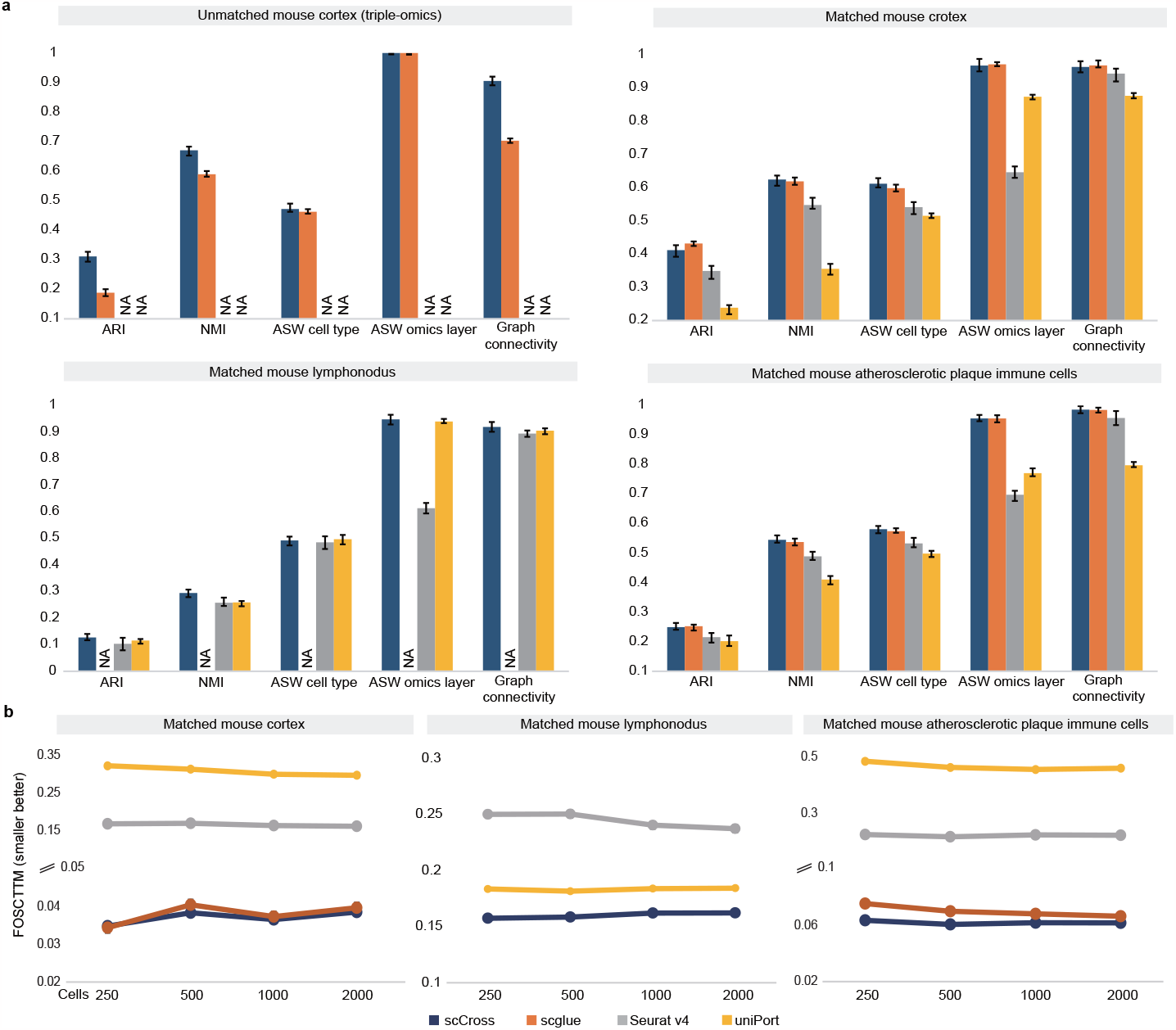
Benchmarking superiority of scCross in single-cell multi-omics data integration. **a**, A comprehensive comparative analysis showcases the standout performance of scCross against other integration methods. Metrics such as average ARI, NMI scores, and ASW for cell types provide insights into clustering quality on the integrated single-cell multi-omics datasets, signifying the retention of biological characteristics after integration. Additionally, the ASW for omics layers and graph connectivity metrics shed light on the effective mixing of different omics post-integration. The datasets studied include the unmatched and matched mouse cortex, matched mouse lymph nodes, and matched mouse atherosclerotic plaque immune cells. “NA” indicates cases where results were not obtainable. **b**, FOSCTTM scores further underscore the exceptional performance of scCross on subsampled datasets of varying sizes, demonstrating its consistent superiority over other integration techniques. This section delves into datasets like the matched mouse cortex, matched mouse lymph nodes, and matched mouse atherosclerotic plaque immune cells.

To further underline our model’s capacity for omics mixing, we assessed the FOSCTTM (Fraction of Samples Closer than the True Match) score [52], a metric quantifying the alignment error in single-cell multi-omics data integration. This score provides insights into how closely related cells from one omics modality are to those from another in the integrated latent space. A lower FOSCTTM score signals a more accurate integration, as the true match is typically closer than many of the other potential matches. On all three matched datasets, scCross consistently achieved the lowest FOSCTTM on all sizes of subsampled datasets, underscoring its alignment capabilities (Fig. 5b). Furthermore, scCross exhibits stable integration performance across an extensive range of hyperparameter settings (Supplementary Fig. S13a-b), significantly reducing the challenge of parameter tuning.

Effective integration is vital for successful downstream activities such as cross-modal generation, multi-omics simulation, and in-silico perturbation. Stellar performance in these subsequent tasks speaks to the exemplary data integration and information retention capabilities of scCross. Notable differences between modalities in the combined latent space can obstruct generation, simulation, or in-silico perturbation, underscoring the necessity of seamlessly merging all modalities into a unified space.

With the advent of high-throughput single-cell technologies, there’s an increasing demand for methods that can effectively process large-scale single-cell multi-omics data, often comprising millions of cells [36, 53]. In our evaluation of scCross’s integrative capabilities, we tapped into over four million human single-cell atlases, consisting of gene expression data for four million cells [54] and chromatin accessibility data for 0.7 million cells [55]. The challenge with such large-scale integration lies not just in the data volume but also in handling extensive heterogeneity, low per-cell coverage, and the imbalances in cell type compositions [36]. Moreover, there’s a common pitfall where existing methods tend to blend minority cell types with the majority, skewing accurate representation and estimation of these pivotal cell groups [56].

scCross addresses these challenges effectively and retains a significant amount of biological data. Unlike existing methods, this approach can differentiate and represent minority cell populations distinctly from the dominant ones. Thanks to the minibatch neural networks [57], scCross achieves sublinear time complexity and roughly linear memory consumption, as highlighted in Supplementary Fig. S13c. Through the strategic use of the generative adversarial network and MNN prior, scCross brings together gene expression and chromatin accessibility data into a comprehensive multi-omics human cell atlas [55], distinguishing the minority cell populations from the majority as demonstrated in Supplementary Fig. S14a-b.

In scglue’s analysis by Cao et al. [36], notable discrepancies between cell type labels across omics modalities were uncovered. Cells labeled as Astrocytes in scATAC-seq data were found to be aligned with Excitatory neuron clusters in scRNA-seq, suggesting a potential mislabeling in the scATAC-seq dataset, which was inferred from biomarker analysis. Furthermore, a cluster identified as Astrocytes/Oligodendrocytes in scATAC-seq was divided and realigned to separate Astrocytes and Oligodendrocytes clusters in scRNA-seq, as indicated by blue cycles in Supplementary Fig. S15a-b. These discrepancies identified by scglue were also replicated by scCross, shown in the blue-highlighted clusters in Supplementary Fig. S14b and Supplementary Fig. S15c. Our analysis further confirms the potential inaccuracy of the Astrocytes annotation in scATAC-seq data, as its top marker genes are not consistent with the biomarkers of Astrocytes in scRNA-seq data (Supplementary Fig. S15d). Both scglue and scCross demonstrate the ability to detect and address potential discrepancies in cell annotations across different omics datasets.

Compared to methods such as scglue [36], scCross demonstrates enhanced capability in identifying minority cell type populations. This is particularly evident in the case of critical minority populations like Extravillous trophoblasts [58–60]—key cells in placental development and function—and Microglia [61–63], which are the primary immune cells in the brain involved in neuroinflammation and tissue repair. These cells are crucial in reproductive and neural systems, respectively. These critical populations are often prone to over-integration with majority cell types in other method like scglue [36] (Supplementary Fig. S15a). However, as highlighted by the red cycle in Fig. S8B, scCross distinguishes them, further validating the integration prowess of our approach. Comprehensive benchmark comparisons between scCross and scglue [36] for this dataset are available in Supplementary Fig. S15e. Alternative approaches such as Seurat v4 and UniPort have not demonstrated their effectiveness in integrating large-scale single-cell multi-omics data with millions of cells.

Given its intrinsic modularity, scCross is not limited to merely a binary omic integration but boasts the capability to consolidate multiple omic layers (e.g., more than 3), attesting to its comprehensive applicability. To showcase this extensive integration potential, we employed scCross on single-cell multiomics data of the adult mouse cortex, encompassing three distinct modalities: gene expression [35], chromatin accessibility, and DNA methylation [16].

In Fig. 6a-d, scCross exemplifies its prowess in aligning the cell types across the three modalities. The alignment is significantly enriched by the incorporation of common genes MNN prior and specific gene sets, thereby ensuring that corresponding cell types from various omic layers converge harmoniously within the latent space. Such alignment not only amplifies the clarity of cell type demarcation across the omic layers but also paves the way for enhanced cell typing efficacy. scCross demonstrates its proficiency in navigating tripleomics datasets, providing a shared cell embedding that effectively distinguishes between cell types for each modality, as depicted with RNA in Fig. 6a. This clear separation is mirrored in the ATAC-seq and snmC-seq modalities, where cell types also form distinct groupings in Fig. 6b and c, respectively. The embedding further shows its strength by closely aligning identical cell types across different modalities, which are visible as clusters in the same regions of the UMAP space. Adding to this, Fig. 6d illustrates the embedding’s capacity for mixing cells from different modalities in the shared space, facilitating the study of cross-modality interactions without the loss of cell-type-specific clustering. This nuanced approach ensures that while cell types from different modalities interweave, they still maintain the fidelity of their distinct embeddings.

**Fig. 6.**
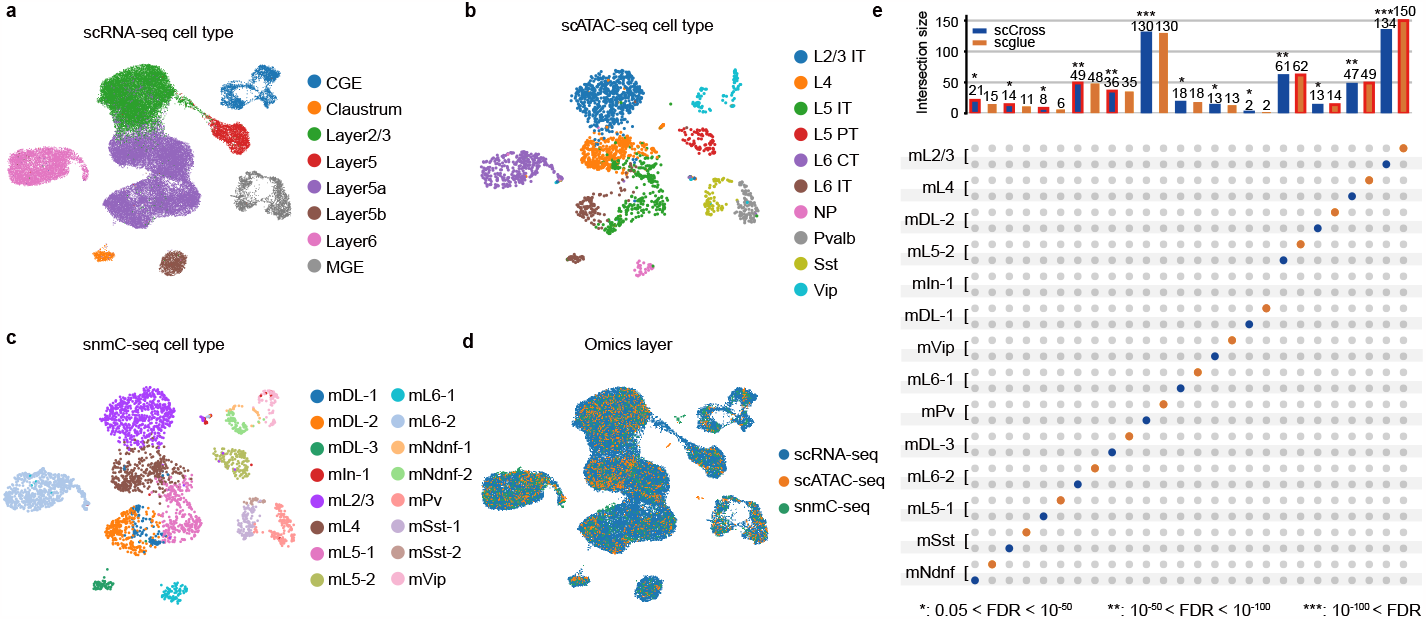
Efficient integration of three omics layers in the mouse cortex by scCross. **a-c**, UMAP visualizations display the aligned cell embeddings, clustering, and cell type annotations for scRNA-seq **a**,, snmC-seq **b**,, and scATAC-seq **c**, on the shared joint latent space, exemplifying the coherence in cellular representations across modalities. **d**, A comprehensive UMAP presentation underlines the flawless intermixing of cells from distinct modalities within the shared latent space, reflecting a true amalgamation of multi-omics information. **e**, An UpSet plot emphasizes the alignment potency of our model, revealing through a three-way biomarker comparison that the distinct modalities are seamlessly integrated within the latent space. Leveraging a three-way Fisher’s exact test [64], our model is showcased to either match or surpass scglue [36] in the alignment of over half of the cell types, with standout results accentuated by a red outline.

Upon establishing the shared cell embeddings, our evaluation centered on the coherence of cell type-specific markers across modalities to validate the alignment accuracy. As depicted in Fig. 6e, there’s a pronounced intersection of markers for each cell type between modalities, substantiating the precision of our cell type alignments. This significant overlap was confirmed by a three-way Fisher’s exact test, with three cell types showing a staggering FDR below 10^*−*100^, and four additional cell types displaying FDRs between 10^*−*100^ and 10^*−*50^. All cell types, with the sole exception of mIn-1, had FDR values indicating strong significance below the 0.05 benchmark. This degree of marker congruence emphasizes the successful integration of the omics layers, as our approach identifies consistent key markers across different modalities for equivalent cell types. In comparison to scglue, our analysis uncovered a greater number of overlapping gene markers in the majority of cell types (9 out of 13), further reinforcing our model’s efficacy in multi-omic data integration.

## Discussion

The ever-evolving domain of single-cell analyses, complemented by multi-omics strategies, continues to redefine our understanding of intricate cellular dynamics. Yet, seamlessly integrating these multi-omics dimensions has remained a challenge. In this context, our scCross method emerges, revolutionizing the way we perceive and amalgamate single-cell multi-omics data. The rigorous testing and evaluations underscore its superior performance, enabling more accurate and comprehensive multi-omics data integration.

But beyond mere integration, scCross introduces the concept of cross-modal generation. This capability allows researchers to derive data from one modality using another, once the model has been trained on a reference single-cell multi-omics dataset encompassing both source and target modalities. Given the often prohibitive cost and experimental challenges of single-cell multi-omics data acquisition, such functionality is crucial. Often, scientists have extensive data in one modality (e.g., RNA-seq) but limited or no profiling in others. With scCross, the knowledge acquired from a comprehensive multiomics dataset can be channeled to generate data in other modalities from a given single modality, filling gaps in our understanding and providing a more complete view of cellular states.

Further accentuating its ability, scCross showcases an array of functionalities tailored for the demands of modern cellular research. Its memory efficiency stands out, ensuring that even datasets with millions of cells are integrated with finesse, making it apt for the vast cell atlases being assembled. Simultaneously, its prowess extends to its capacity for single-cell multi-omics data simulation. Through judicious sampling from the joint latent space and adjusting omic-decoders, it can simulate matched single-cell datasets across various modalities for the same consistent set of cells. This serves not only for data augmentation but also as a benchmarking tool, given the known ground truth in these simulated datasets. Building on these capabilities, scCross offers the powerful in-silico perturbation functionality. As evidenced in cases like COVID-19 cellular dynamics, it offers a fresh perspective to formulate and probe potential intervention strategies to modulate cellular states. Once primed with a reference multi-omics dataset, it can be harnessed on single-modality data, enabling intricate multi-omic in-silico explorations.

The scCross method presents significant potential for the single-cell research community. Its distinct functionalities and reliable performance position it as a potentially valuable tool for researchers in the field of single-cell multi-omics analyses. scCross facilitates the integration of different modalities, supports comprehensive data generation, and enables detailed simulations and perturbations. These capabilities could contribute substantially to advancing the study of complex biological systems.

## Conclusions

scCross has the potential to significantly influence the field of single-cell multiomics research. Its ability to bridge modalities effectively, along with capabilities in cross-modal generation, multi-omics simulation, and augmentation, may enhance both the speed and depth of biomedical research. Additionally, its suitability for in-silico explorations offers practical applications beyond academic research, potentially assisting in the design of targeted experiments for cellular state manipulations and possibly hastening the translation of biomedical research into real-world applications.

## Methods

### Single-cell data processing

The single-cell data employed in our study follows specific preprocessing pipelines tailored to each modality. Initial procedures, such as cell calling, quality control, and other relevant steps, can follow standard single-cell data processing methods and are often performed by the original data sources [17, 18, 31, 35]. Within our workflow, these matrices are further transformed to produce cell vectors optimized for neural network input. Besides, these matrices’ feature sets are denoted as 𝒮_*k*_, where *k* ranges from 1 to *K. K* symbolizes the number of distinct omics data types collected. For example, in scRNA-seq, *𝒮*_*k*_ represents a gene list, while in scATAC-seq, it represents a set of chromatin regions. Profiling matrices from the *k*^*th*^ modality is represented as *χ*_*k*_. Individual cells from this modality are given by 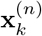, and *N*_*k*_ indicates the sample size.

For single-cell transcriptomic data, specifically scRNA-seq, we adopt Scanpy [9] (v.1.8.2). The cell-by-gene expression matrix experiences normalization, log-transformation, and scaling. Dimensionality reduction follows, with PCA, as realized in Scanpy [65], being our method of choice. On the epigenetic data front, modalities like scATAC-seq and snmC-seq demand distinct processes. SnmC-seq data transitions through a sequence of normalization, log-transformation, scaling, and PCA, utilizing epiScanpy [66] (v.0.4.0). Given the inherent sparsity of the scATAC-seq data matrix, dimensionality reduction employs the LSI (Latent Semantic Indexing) function of scCross, grounded in the latent semantic indexing algorithm [14, 67].

To facilitate the calculations of gene sets matrix and common gene MNN priors, it is crucial to have a consistent gene-centric representation across different modalities. Due to the inherent nature of MNN priors, the single-cell data from diverse modalities needs to be translated into activities associated with genes. To achieve this uniform representation, we employ the geneactivity function in epiScanpy [66] (v.0.4.0), transforming various omics data types (like scATAC-seq and snmC-seq) into gene activity. The gene formed data (Gene expression/Gene activity score) of each modality *k* is represented with 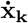. It is obvious that 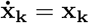 when *k*^*th*^ modality is scRNA-seq.

It’s noteworthy that our model’s applicability isn’t confined to the discussed modalities. For any modality, the transformation of each cell into a real-valued vector is a must. Often, normalization is required to address technical nuances, such as variations in library size among cells. For modalities not tackled in this work, we advocate adhering to the modality-specific standard single-cell preprocessing protocol when generating the input vectors.

### Curating gene sets from Gene Ontology and Pathway terms

In our study, gene sets 𝒢 𝒮 are tapped as an avenue to integrate functional biological priors, enhancing the representation learning of cells across different modalities. This, in turn, bolsters the effectiveness of single-cell data integration and generation efforts, as these tasks are fundamentally rooted in effective cell embeddings.

We utilize a rich collection of gene sets, comprising 7,481 sets from the GO Biological Process ontology (denoted as c5.go.bp in MSigDB [68]), 2,922 gene sets curated from pathway databases (c2.cp in MSigDB [68]), and 335 sets of transcription factor targets sourced from [69]. Gene sets related to biological processes bear the prefix “GOBP”, while those derived from pathway databases are assigned prefixes echoing their respective sources, such as KEGG [70], WP [71], and REACTOME [72].

To effectively harness the gene sets, 𝒢 𝒮, each gene set is characterized based on the genes it contains. In the context of scRNA-seq data, a gene set is represented by the expression levels of its constituent genes. This provides a direct quantification of the gene set’s activity within a cell, crucial for grasping cellular states and functions. However, with non-transcriptomic modalities, such as scATAC-seq, where direct gene expression values are absent, we resort to the geneactivity function from epiScanpy [66] (v.0.4.0). This tool calculates a gene activity matrix, serving as an effective surrogate for gene expression. Using this matrix, we can obtain the binary matrix 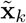, which is the correlation of genes in gene expression/activity matrix and genesets in the databases we mentioned above. We utilize the approach of [73] where we set the correlation of a gene to a geneset to 1 if the gene is contained in the geneset, 0 otherwise. Then, we obtain the cell by geneset matrix with the function:

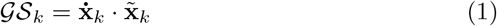

where 𝒢 𝒮_*k*_ means the cell by geneset feature matrix of modality *k*. These scores, once extracted, are integrated with the processed input features, grounding our analyses in solid biological contexts.

### Deep generative neural network structure

To learn cell embeddings from omics data, we employ a variational autoencoder (VAE).

Our model initially projects **x**_*k*_ into a low-dimensional latent space, **z**_*k*_, utilizing a variational approach. The encoder function is formulated as:

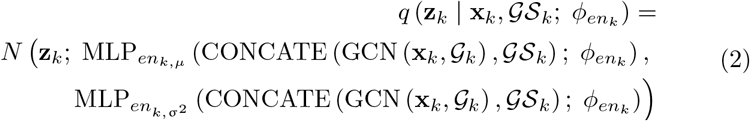

Here, 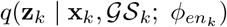 describes the encoder part of the VAE for the *k*^*th*^ modality, CONCATE means concatenating two matrices together. It uses a graph convolutional network (GCN) to model the latent space **z**_*k*_ given the observed data **x**_*k*_.

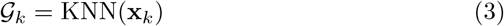

The graph 𝒢_*k*_ is constructed using the k-nearest-neighbor algorithm and serves as the adjacency matrix for the GCN.

The data likelihoods *p* (**x**_*k*_ | **z**_*k*_; *θ*_*k*_), which represent the data decoders, are constructed using a multilayer perceptron. The specific formulation of the data likelihood depends on the distribution of the omics data. In the case of count-based scRNA-seq and scATAC-seq data, our model utilizes the negative binomial (NB) distribution. The NB distribution is defined as follows:

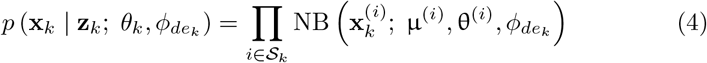

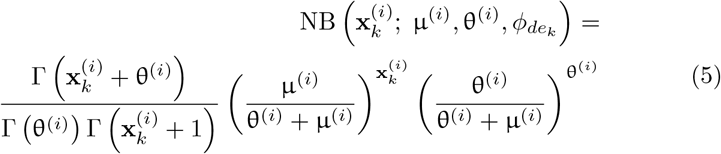

In the equation above, 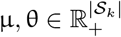 are the mean and dispersion of the negative binomial distribution. Additionally, 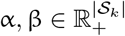 are scaling and bias factors. The Hadamard product is denoted by ⊙, Softmax^(i)^ represents the *i*^*th*^ dimension of the softmax output, and 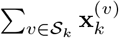 provides the total count in the cell ensuring the library size of reconstructed data matches the original.

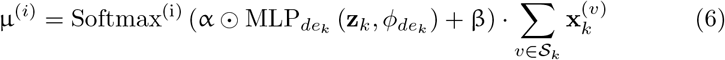

The set of learnable parameters in this context is *θ*_*k*_ = *{*α, β, θ*}*. Mean-while, 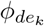 denotes the set of learnable parameters in the multilayer perceptron decoder of the *k*^*th*^ omics layer, 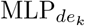. The flexibility of this model also allows for the incorporation of many other distributions.

Finally, the optimization process aims to maximize the evidence lower bound:

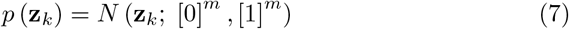

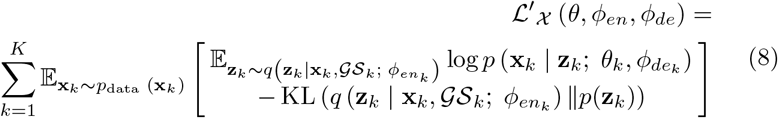

This equation represents the loss function that the VAE optimizes, where *m* means the length of latent space vector which is 50 and 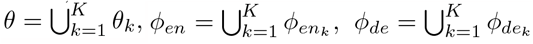, are the combined sets of encoder and decoder parameters, respectively. It incorporates both the data likelihood and a Kullback-Leibler (KL) divergence term to enforce a meaningful latent space for each of the *K* modalities.

### Multi-omics data integration, crossmodal generation, and simulation

To harmonize the latent embeddings produced from different omics modalities, we implement a cross-modal bidirectional alignment method utilizing multi-layer perceptrons (MLPs):

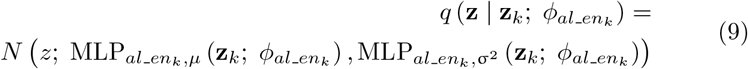

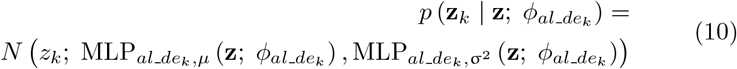

In these equations, 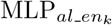 denotes the aligner encoder, which converts the latent embedding of the *k*^*th*^ omics layer to a collective latent space. On the other hand, 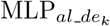 represents the aligner decoder, which translates the unified latent space embedding back to the latent space of the *k*^*th*^ omics modality. This alignment process is key to converting single-cell multi-omics data into a shared latent representation, denoted as **z**. By establishing such a common latent space, we facilitate an effective integration of disparate omics data.

The architecture of scCross operates on a two-step variational autoencoder (VAE) framework to encode omics layers into a merged space. Inverting this methodology, any encoded data in this unified space can be reverted to any particular omics layer’s latent representation using a dual-step decoding procedure. The benefit here is the cost-efficiency: rather than requiring *k*! distinct encoder/decoder combinations to synchronize any two modalities, this method necessitates only a couple. This streamlined approach facilitates multi-omics simulations across any number of omics layers. Nevertheless, the method’s success heavily depends on the precise integration of data in the shared latent space. If discrepancies emerge in this unified space between modalities, the generated data will mirror those differences.

By leveraging a bi-directional aligner, the cross-modal generation of data from modality *j* to modality *h* can be conceptualized with the following equation:

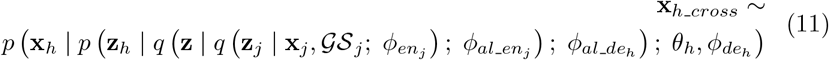

where *x*_*h cross*_ means cross generated *x*_*h*_ from *x*_*j*_. A visual representation of the described procedure is illustrated in Supplementary Fig. S1a. The process begins with the input vector from modality *j*, which undergoes dimensionality reduction and mapping to the joint latent space *z*. Subsequently, the decoder associated with modality *h* is employed to reconstruct *z* into the data vector for modality *h*.

Notably, when *j* is not equal to *h*, this process facilitates cross-modal generation, allowing the transformation of information from one modality to another. Conversely, when *j* is equal to *h*, it becomes a mechanism for self-augmentation within the same modality. A visual representation of the described procedure is illustrated in Supplementary Fig. S1b.

Similarly, single-cell data simulations tailored to specific cellular states for a particular modality *h* are described as:

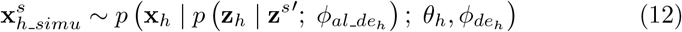

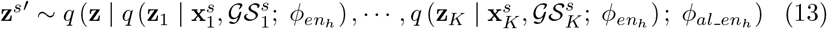

In this context, 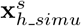 means simulated data of *h* modality for specific cellular states, 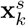 pertains to the feature matrices of specific cellular states for *k*^*th*^ modality, 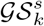 is the gene set matrices for *k*^*th*^ modality of specific cellular states, 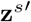 represents the sample result of 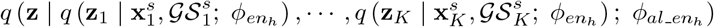. simulate single-cell multi-omics for a certain cell population. We first sample in the joint latent space to produce latent embedding vectors to represent cells from the specific cell population. The number of samples determines the number of simulated cells. The latent embedding *z*_*h*_ will then be converted back into the input vector *x*_*h*_ via specific modality-specific decoder. A visual representation of the described procedure is illustrated in Supplementary Fig. S1c.

The model’s fitting procedure seeks to maximize the evidence lower bound expressed as:

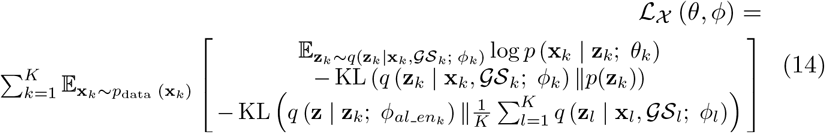

Here, the set of encoder and decoder parameters are expressed as 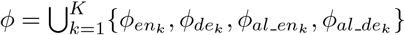 and 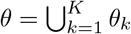, respectively.

### High-quality data integration and generation via generative adversarial learning

In the endeavor to ensure precise alignment of diverse omics data, we harness the potential of a generative adversarial learning strategy, a method that showcases its prowess in numerous prior studies [36, 74].

Central to our model is a discriminator, denoted as D_*z*_, equipped with a *K*-dimensional softmax output, designed specifically for the seamless alignment of various omics layers [35, 75]. Functioning based on the latent space embeddings of cells, denoted as *z*, D_*z*_ endeavors to identify the omics layer origin of the cells. The training of this discriminator is steered by the minimization of the multi-class classification cross entropy:

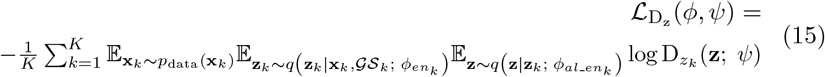

Here, 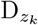 symbolizes the *k*th dimension of the discriminator output, while *ψ* refers to the suite of parameters, amenable to learning, within the discriminator. Contrarily, the data encoders are trained with the ambition to deceive this discriminator, a strategic move to fortify the alignment of cell embeddings stemming from distinct omics layers.

Furthermore, to bolster the generation of cross-modality data, we present unique discriminators for each modality. This approach aids in refining the reconstruction of the data:

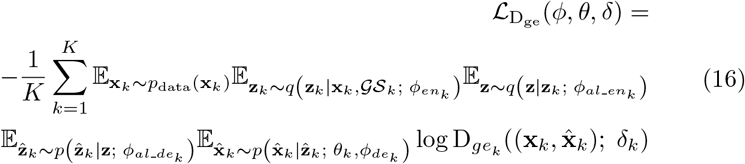

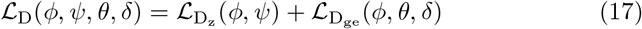

Within this context, 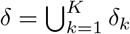 encapsulates the collection of learnable parameters in the discriminators. Additionally, 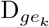 represents the output from the *k*^*th*^ discriminator, tailored to reinforce the congruence between the model-generated data 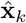 and the real data **x**_*k*_ for the *k*^*th*^ omics layers. The terms 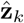 and 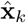 depict the data generated by the aligner decoder and VAE decoder for the *k*^*th*^ modality, respectively.

### Common gene MNN prior to align cells in similar cellular states

For an effective alignment across various modalities, it is essential to harness common genes that span across modalities, such as gene expression in scRNA-seq and gene activity in scATAC-seq and snmC-seq. These genes serve as a pivotal anchor, ensuring consistency and coherence across modalities. While previous omics data integration methods have been successful in achieving a semblance of agreement in the data distribution, the main challenge remains. The true test of alignment is whether embeddings of the same or similar cells across different modalities can be represented identically or similarly within the shared latent space.

Here, the effective multi-modal alignment is achieved by utilizing the Mutual Nearest Neighbors (MNN) approach, which has been shown to effectively align cells of analogous cellular states across different omics data [76]. First, each modality is represented by a matrix, 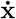, comprised of cells by common genes. Subsequently, the MNN-corrected common genes matrix from each modality is processed using PCA. This produces a common gene prior, *G*, which encapsulates all modalities:

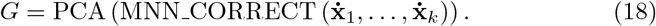

The alignment loss function, ℒ_MNN_, is then formulated to minimize the difference between the cosine similarities of the latent embeddings and their corresponding common gene priors:

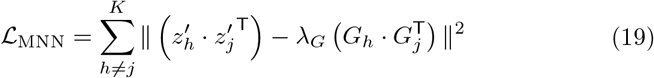

where 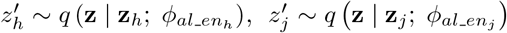 and *λ*_*G*_ are the cell embedding in the shared space for the modality h and j, respectively, generally set to 1, assists in scaling the difference between the two subtractions. In this equation, *G*_*h*_ and *G*_*j*_ refer to the common gene priors for modality *h* and *j* corresponding to the cells’ embedding 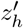 and 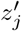, respectively.

### Overall training strategy

The training process for scCross unfolds over two distinct stages. Initially, the primary objective is the isolated training of a Variational Autoencoder (VAE) for each modality. This approach enables us to capture and understand the unique biological information inherent within each modality before proceeding to their integration. The loss function for this first stage is represented as:

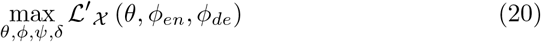

Transitioning to the second stage, the trained VAE models from the preliminary stage serve as a foundation. At this juncture, bi-directional aligners are incorporated, merging the modalities into a unified latent space. Discriminators also come into play during this stage, contributing to the integration of multi-omics data. The associated training losses for this stage are detailed as:

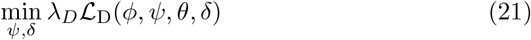

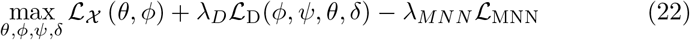

In these equations, the terms *λ*_*D*_ and *λ*_*MNN*_ dictate the relative contributions of the discriminator and the common gene MNN prior, respectively. To optimize the training of the scCross model, we utilize the stochastic gradient descent technique. Notably, the RMSprop optimizer, devoid of a momentum term, is selected to enhance stability during adversarial training.

### Cross-modal generation of single-cell data

Cross-modal generation of single-cell data serves diverse application scenarios. Primarily, our methodology focuses on training our model using a reference multi-omics dataset encompassing both source and target modalities. Subsequently, this trained model facilitates the generation of the missing modality in a untrained dataset. An alternative application stems from cases with limited data for one modality. To demonstrate our method’s prowess, we train it on such partial datasets and leverage this for cross-modal generation.

Initially, we employ the unmatched mouse cortex dataset [35] to train our model. Post this preliminary training, we execute fine-tuning using the scATAC-seq component of the matched brain dataset [31]. We then project the scATAC-seq data from this matched dataset to its corresponding scRNA-seq, using the latter’s original data as a benchmark. To showcase the capability of our model in handling datasets with limited modalities, we design a simulation using the matched mouse cortex dataset [31]. The dataset is bifurcated: 20% is employed for model training, and the remaining serves for cross-generation and validation. This simulation essentially aims at highlighting how our model, when trained on single-cell multi-omics data with a restricted modality, can upscale and extract valuable insights from the limited data. After the training phase, we employ our model to cross-generate the scRNA-seq data from the available scATAC-seq modality, thereby demonstrating the model’s ability to effectively leverage the trained insights from the limited dataset.

The robustness of our cross-modal generation technique is underscored by a comprehensive downstream analysis. The generated data is first clustered using the Leiden algorithm [77]. With the assistance of Scanpy [9], we discern the top 100 differentially expressed genes (DEgenes) for each cell type in both the generated and original scRNA-seq datasets. These DEgenes not only facilitate the transfer of cell type annotations from the original to the generated datasets but also serve as potential biomarker genes. Cell types with the majority of shared genes are aligned. Pathway enrichment further enriches our validation process. For each cell type, pathway associations rooted in the top 100 DEgenes are pinpointed via the ToppGene portal [34]. Concurrently, to decipher the intricate interplay between cells, CellChat [78] is employed for detailed cell-cell interaction analysis. To gauge the accuracy and resemblance between our generated and the original scRNA-seq data, we deploy an array of validation metrics. Pearson correlation [79] coefficients provide insights into decomposition, cell-cell interactions, and pathway analyses.

### Single cell multi-omic data simulation

Our model’s training is grounded on the matched mouse cortex dataset [31]. Specifically, for our simulation exercise, we choose the cellular type Ast as the target cellular status. Leveraging our model, we generate data sets at one-fold (1X) and five-fold (5X) of the original count of Ast cells’ single-cell multiomics data. These data sets are subsequently visualized using both Scanpy [9] (v.1.8.2) and epiScanpy [66].

With the assistance of Scanpy [9], we discern the top 100 differentially expressed genes (DEgenes) for Ast cells in both the 5X simulated and original scRNA-seq data. GO and pathway enrichment in both data further enriches our validation process. GO and pathway associations rooted in the top 100 DEgenes are pinpointed via the ToppGene portal [34].

To assess the level of correlation between the differential gene signatures of the original Ast cells and the simulated 1X Ast data against a background, we employ the RRHO methodology [80].

To further harness and validate the simulated 1X Ast single-cell multiomics data, we input it into various integration techniques, notably our scCross and the externally established scglue [36]. The outcomes from these integrative analyses serve to demonstrate the adeptness of the scCross model in generating high-quality matched single-cell multi-omics data for specific cellular states, which can be potentially leveraged to benchmark other single-cell multi-omics data integration methods.

### In-silico perturbations and explorations

In this section, we aim to showcase the in-silico perturbation pipeline of scCross, using the single-cell multi-omics dataset of COVID-19 as an exemplar. This dataset, comprising both scRNA-seq and ADT data (CITE-Seq), serves as a suitable backdrop for our demonstration, given its comprehensive nature that facilitates deeper insights into the disease [17]. The preliminary step in our pipeline involves visualizing the scRNA-seq data of the COVID-19 dataset. We utilized the Scanpy package for this purpose [9] (v.1.8.2), generating a UMAP plot depicted in Fig. 4. Concurrently, the same package aided in the identification of the highly variable genes. With these genes in focus, the scCross model, once trained on matched scRNA-seq and single-cell protein datasets [17], embarked on the in-silico perturbation process. Each highly variable gene underwent a virtual perturbation on COVID-19 patient cells, and the model gauged and scored its influence on bridging the latent space gap between contrasting disease states, namely healthy and COVID-19 conditions. The culmination of this process results in a ranked list of genes, with top scorers spotlighted as potential RNA bio-markers for COVID-19.

To further visualize the ramifications of these perturbations, we employed the Seaborn Python package [81] (v.0.11.2). This involved computing the cosine distances in the latent space between COVID-19 and healthy cells during both upward and downward perturbations (scaled by *±*1 in log space) of each gene. The resultant visualization, available in Fig. 4b, exclusively displays genes that effectively narrowed the latent space divide.

To further analyze the genes obtained from perturbation, we perform pathway analysis using the ToppGene website [34] with the down-perturbed genes that make the latent space closer. Additionally, we use the t-test [82] and Wilcoxon [83] tests, which are general methods for differential gene expression analysis, to generate the same number of DEgenes as our approach between COVID-19 patient cells and healthy cells. The pathway analysis is also performed using the ToppGene website [34]. We select several pathways highly associated with COVID-19 to demonstrate the effectiveness of our approach in identifying disease biomarkers via in-silico perturbations (Fig. 4c).

Furthermore, we execute cross-generation of single-cell protein data using the scRNA-seq dataset. For this procedure, we perturb the relevant genes identified in the prior analysis within the healthy scRNA-seq data. This gene perturbation leads to the creation of perturbed COVID-19 scRNA-seq data, which is subsequently harnessed for cross-generation in order to produce a generated dataset representing single-cell COVID-19 protein data.

To evaluate the correlation between the similarity of differential protein expressions in the original COVID-19 protein data and the perturbed, generated COVID-19 protein data against the backdrop of healthy protein data, we employ RRHO [80] as a visualization technique.

In order to validate the degree of resemblance between the up-perturbed and cross-generated single-cell protein data and the original dataset, we leverage the Scanpy package [9] (v.1.8.2). This comparison is performed within a comprehensive landscape encompassing all cells (Fig. 4f). Pearson correlation [79], along with its associated *p*-values, is adopted to quantitatively affirm the similarity between these datasets.

### Single cell multi-omics data integration

We evaluate the agreement between the cell-type labels and the Leiden algorithm [77] clusters obtained from the integrated dataset for each omic separately using Normalized Mutual Information (NMI). NMI measures the overlap between two clusterings, with scores ranging from 0 (no overlap) to 1 (perfect agreement). We perform the Leiden clustering [77] with a resolution of 1 for all methods and datasets. We use the scikit-learn [84] (v.0.22.1) implementation of NMI.

The Rand index is another metric that compares the overlap of two clusterings. We compare the cell-type labels with the Leiden clustering [77] computed on the integrated dataset for each omic separately. An ARI of 0 or 1 corresponds to uncorrelated clustering or a perfect match, respectively. We perform the Leiden clustering [77] with a resolution of 1 for all methods and datasets. We use the scikit-learn [84] (v.0.22.1) implementation of the ARI.

In our work, there are two kinds of ASW. One is cell type ASW, another is omics layer ASW. Cell type ASW is used to evaluate the cell type resolution. It can metrix model’s clustering ability. Cell typ ASW is defined as in a recent benchmark study [36, 51]:

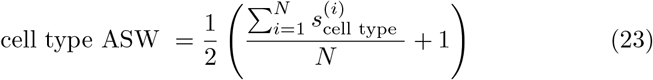

where 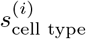 is the cell type silhouette width for the *i*^*th*^ cell, and *N* is the total number of cells. Cell type ASW has a range of 0 to 1. Higher values indicate better cell type clustering resolution.

Omics layer ASW is used to evaluate the integration mixing ability among omics layers. It is defined as in a recent benchmark study[36, 51]:

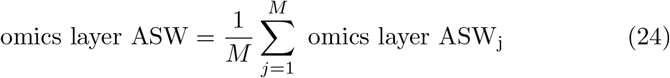

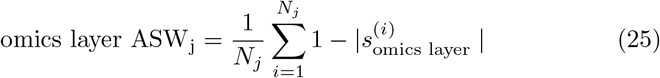

where 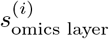 is the omics layer silhouette width for the *i*^*th*^ cell, *N*_*j*_ is the number of cells in cell type *j*, and *M* is the total number of cell types. Omics layer ASW has a range of 0 to 1. Higher values indicate better integration mixing ability.

The graph connectivity metric [36, 51] is used to estimate the ability to mixing different omics data:

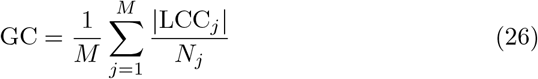

where LCC_*j*_ is the number of cells in largest connected component of the cell k-nearest neighbors graph (K = 15) for cell type *j, N*_*j*_ is the number of cells in cell type *j* and *M* is the total number of cell types. Graph connectivity has a range of 0 to 1. Higher values of graph connectivity indicate better mixing ability.

The FOSCTTM metric [36, 52] is used to evaluate the single-cell level alignment accuracy. It is computed on two datasets with known cell-to-cell pairings. Suppose that each dataset contains *N* cells, and that the cells are sorted in the same order, that is, the *i*^*th*^ cell in the first dataset is paired with the *i*^*th*^ cell in the second dataset. Denote *x* and *y* as the cell embeddings of the first and second dataset, respectively. The FOSCTTM is then defined as:

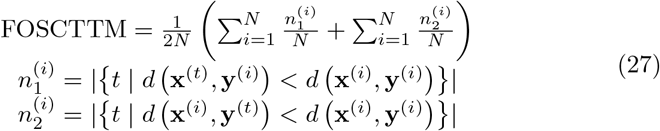

where 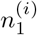 and 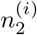 represent the number of cells in the first and second dataset that are closer to the *i*^*th*^ cell than their true matches in the opposite dataset. The Euclidean distance, denoted as *d*, is used to calculate the distance between cells. FOSCTTM ranges from 0 to 1, and lower values indicate higher accuracy.

To train the atlas level large scale data, we made several adaptations to our training process. We employed the Leiden algorithm [77] to cluster the original scRNA-seq and scATAC-seq data and obtained the cluster numbers. Then, we aggregated and averaged the cells within each cluster to create a meta cell representing that cluster. The common gene MNN prior for cells within each cluster was calculated based on these meta cells. To boost training efficiency, we utilized the MNN loss, calculated based on the meta cells instead of all cells, for the second stage of fine-tuning scCross (loss defined in eqn.22). Finally, we utilized Scanpy [9] (v.1.8.2) to generate the UMAP visualization.

In the triple omics integration, as the snmC-seq data’s distribution is close to zero-inflated log-normal (ZILN) distribution, we use that distribution in our data decoder:

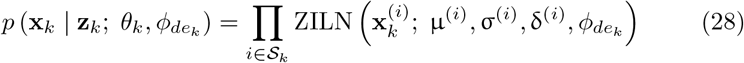

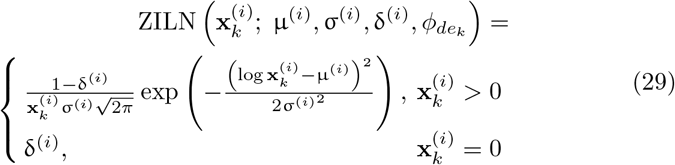

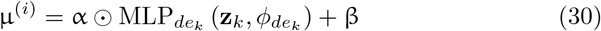

In the equation above, 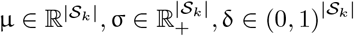 are the log-scale mean, log-scale standard deviation and zero-inflation parameters of the zero-inflated log-normal distribution, respectively. Additionally, 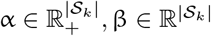 are scaling and bias factors. The set of learnable parameters in this context is *θ*_*k*_ = {σ, δ, α, β} and 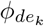.

To further verify our model’s triple omics integration ability, we employ a nearest neighbor-based label transfer approach using the ATAC-seq dataset as the reference and test the cell similarity of cells with the same label in different omics by markers. For each cell in the scRNA-seq and snmC-seq datasets, we identify the five closest neighbors in the ATAC-seq dataset within the shared embedding space. We achieve label assignment through a majority voting system based on these neighbors. To confirm the accuracy of our alignment, we test for significant overlap in cell type marker genes across modalities. Initially, we convert features from all omics layers into gene-centric representations with geneactivity function in epiScanpy [66]. Subsequently, we pinpoint cell type markers for each omics layer. We adopt a one-versus-rest Wilcoxon rank-sum test to ascertain these markers, using a significance threshold of FDR *<* 0.05. The significance of overlapping markers among the three omics datasets is gauged using the three-way Fisher’s exact test [64]. Visualization of the significant markers, as well as a comparison of FDR with the scglue method [36], is facilitated by the UpSet plot [85].

### Batch effect correction

Batch effect intro modalities can be a hard problem to resolve when integrating multi-modal datasets. Assuming *b ∈* 1, 2, …, *B* is batch index, where *B* is the total number of batches in all modalities. Here, we utilize the batch effect correction method in [36], which is introduce batch-specific parameters in VAE data decoder:

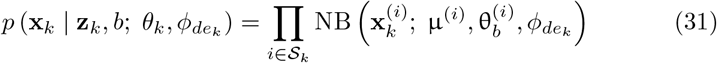

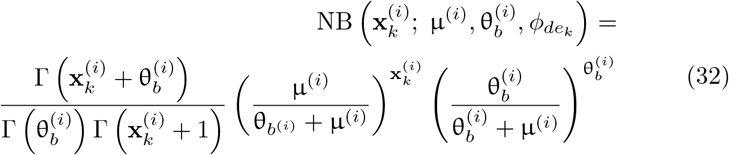

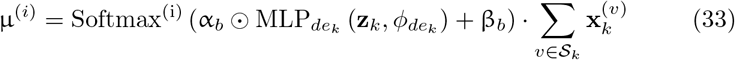

where 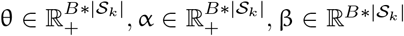, and θ_*b*_, α_*b*_, β_*b*_ are the *b*^*th*^ row of θ, α, β. Other VAE decoders in different distribution can also be extended in similar ways.

### Implementation details

In our model, we employ linear dimensionality reduction techniques such as PCA (Principal Component Analysis) [65] for scRNA-seq data, gene information, and gene set information. For scATAC-seq data, we use LSI (Latent Semantic Indexing) [67] as the first transformation layers of the data encoders (while the decoders were still fitted in the original feature spaces). These canonical methods effectively reduce the model size and allow for modular input, enabling the use of advanced dimensionality reduction or batch effect correction methods as preprocessing steps for scCross integration.

To balance the losses in our model, we determine the value of lambda (*λ*) for loss balancing. The lambda selection process can be found in Supplementary Fig. S13a-b (please refer to the supplementary materials of the corresponding publication).

Our model consists of two steps of training. In the first step, each omic layer’s VAE is trained separately for each modality. This step aims to obtain well-preserved bio-information in the latent embeddings (represented as **z**_*k*_) without interference from other modalities. In the second step, integration is performed by training all modalities’ **z**_*k*_ together to achieve a unified latent space represented as **z**. This step enables the integration of different omic layers in the scCross model. The set of the best hyperparameters is presented in Supplementary Fig. S13a-b.

## Data availability

The datasets used in this study have been obtained from various sources. The matched mouse cortex dataset [31] is obtained from Gene Expression Omnibus (GEO) repository under the following accession numbers GSE126074. The matched mouse atherosclerotic plaque immune cells dataset is got from Gene Expression Omnibus (GEO) repository under the following accession numbers GSE240753. The matched mouse lymphonodus dataset [50] is downloaded from Gene Expression Omnibus (GEO) repository under the following accession numbers GSE140203. The unmatched mouse cortex scRNA-seq dataset [35] is publicly available in the website at dropviz.org. The unmatched mouse cortex snmC-seq dataset [18] is publicly available and obtained from the Gene Expression Omnibus (GEO) repository under the following accession numbers GSE97179. The unmatched mouse cortex scATAC-seq dataset is downloaded from the 10X Genomics website at support.10xgenomics.com. The human cell atlas dataset [54, 55] is obtained from Gene Expression Omnibus (GEO) repository under the following accession numbers GSE156793 for scRNA-seq and GSE149683 for scATAC-seq. Lastly, the COVID-19 dataset [17] used in the study is obtained from the website at covid19.cog.sanger.ac.uk. Detailed average benchmarking information for our integration evaluation across methods section is available in Supplementary Data S1.

## Code availability

The scCross framework is implemented in the “scCross” Python package, which is available at https://github.com/mcgilldinglab/scCross.

## Acknowledgments

This work is partially supported by grants awarded to JD from the Canadian Institutes of Health Research (CIHR) (Grant No. PJT-180505), the Fonds de recherche du Québec - Santé (FRQS) (Grant Nos. 295298 & 295299), and the Natural Sciences and Engineering Research Council of Canada (NSERC) (Grant No. RGPIN2022-04399), along with support from the Meakins-Christie Chair in Respiratory Research. The study also benefits from partial funding to HW through grants from the National Natural Science Foundation of China (Grant Nos. 62272278 & 61972322), the National Key Research and Development Program (Grant No. 2021YFF0704103), and the Fundamental Research Funds of Shandong University. Additionally, KKM has received partial support for this research from the CIHR (Grant No. PJT-166142).

## Author contributions

JD and XY conceived the study; JD, XY and HW designed the analyses and experiments; XY performed the analyses, experiments and visualizations; XY and JD prepared the first draft; JD, XY, HW and KKM reviewed and edited the draft; KKM contributed the data GSE240753.

