## Supplementary Figs. S1-S15 for "scCross: A Deep Generative Model for Unifying Single-cell Multi-omics with Seamless Integration, Cross-modal Generation, and In-silico Exploration"

Yang et al.

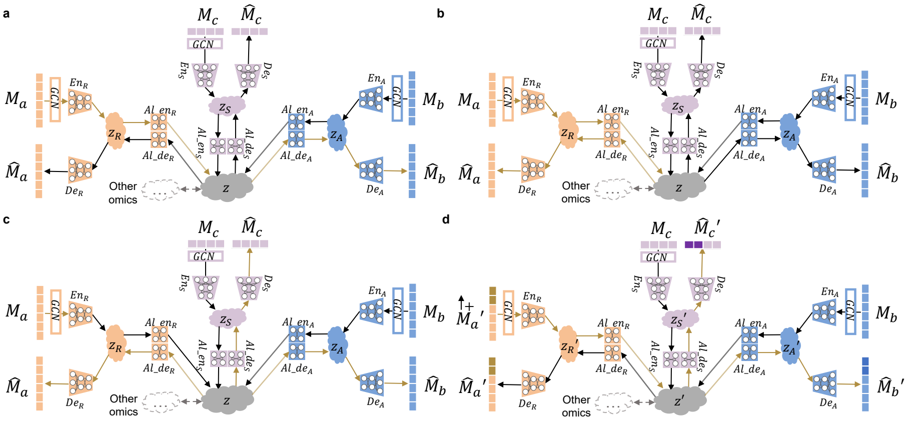

**Supplementary Fig. S1 Data flow of the scCross functions. a,** Cross-Generation Function Data Flow: Data from modality a is encoded into a common latent space z using its specific encoder and then decoded into modality b with modality b's decoder. **b,** Self-Enhancing Function Data Flow: Data from modality a is encoded into the common latent space z through its encoder and then decoded back into modality a itself with its own decoder. **c,** Multi-Omics Simulation Function Data Flow: The mean latent space distribution of a specific cluster of cells is sampled and decoded into each data type, forming a multi-omics simulation. **d,** Perturbation Function Data Flow: Several features in modality a are perturbed, encoded into the common latent space, and then other data types with the same perturbation can be decoded from the latent space using their own decoders.

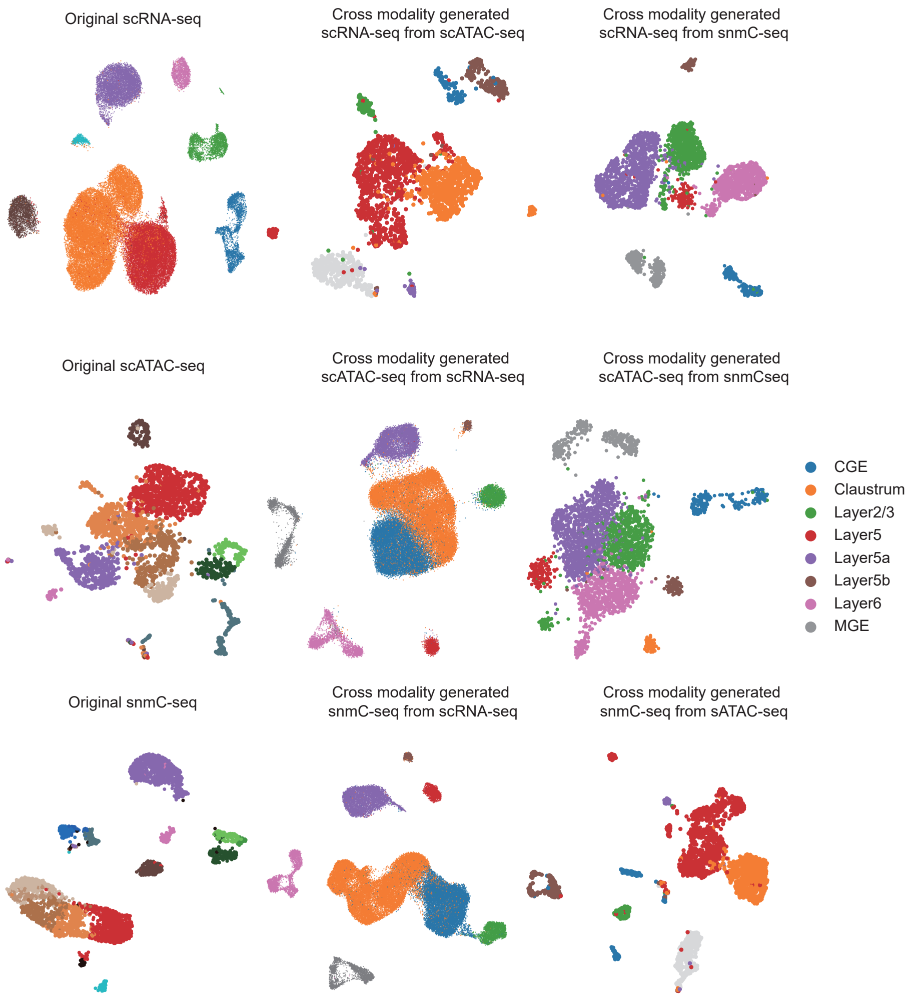


**Supplementary Fig. S2 UMAP visualization of the original and cross generated data in unmatched mouse cortex dataset.** The UMAP visualization compares the original scRNA-seq and scATAC-seq data, along with the cross-generated data for each other in this unmatched mouse cortex dataset.. Notably, the cross-generated data showcases a substantial similarity with the original data in UMAP, emphasizing the effectiveness of the integration process in this dataset.

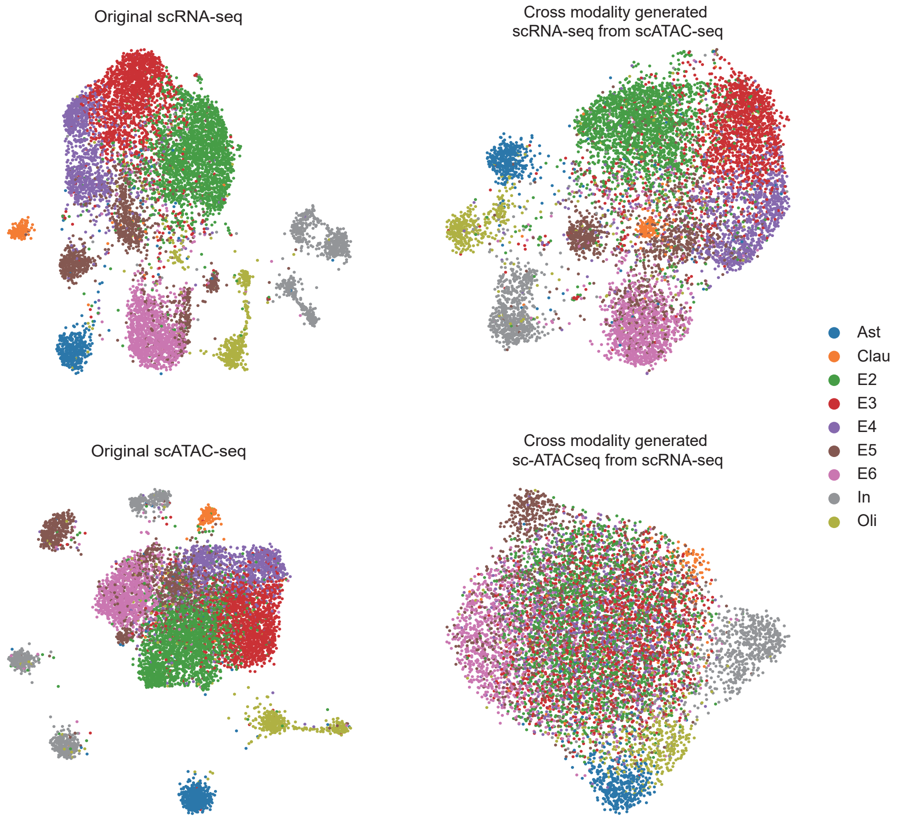


**Supplementary Fig. S3 UMAP visualization of the original and cross generated data in matched mouse cortex dataset.** UMAP visualization comparing the original scRNA-seq and scATAC-seq data of matched mouse cortex, as well as the cross-generated data for each other. The UMAP visualization clearly illustrates that the cross-generated data exhibits a high degree of similarity with the original data in the matched mouse cortex dataset.

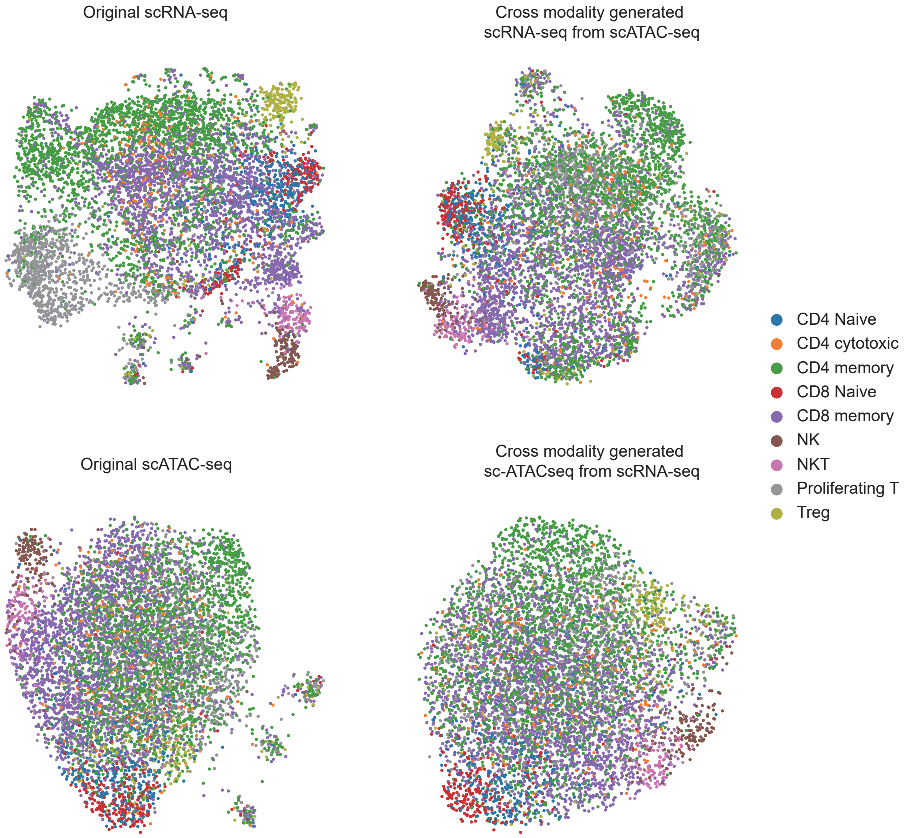


**Supplementary Fig. S4 UMAP visualization of the original and cross generated data in matched mouse lymphonodus dataset.** UMAP visualization comparing the original scRNA-seq and scATAC-seq data of matched mouse lymphonodus, as well as the cross-generated data for each other. Remarkably, the cross-generated data displays a high degree of similarity with the original data in UMAP within the matched mouse lymphonodus dataset.

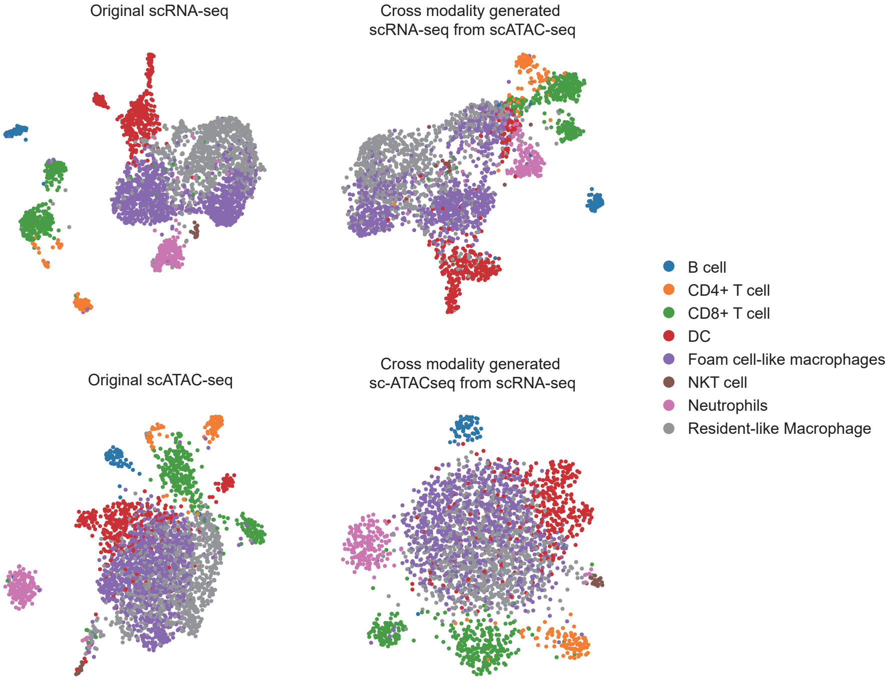


**Supplementary Fig. S5 UMAP visualization of the original and cross generated data in matched mouse atherosclerotic plaque immune cells dataset.** UMAP visualization comparing the original scRNA-seq and scATAC-seq data of matched mouse atherosclerotic plaque immune cells, as well as the cross-generated data for each other. Impressively, the cross-generated data exhibits a high degree of similarity with the original data in UMAP within the matched mouse atherosclerotic plaque immune cells dataset.

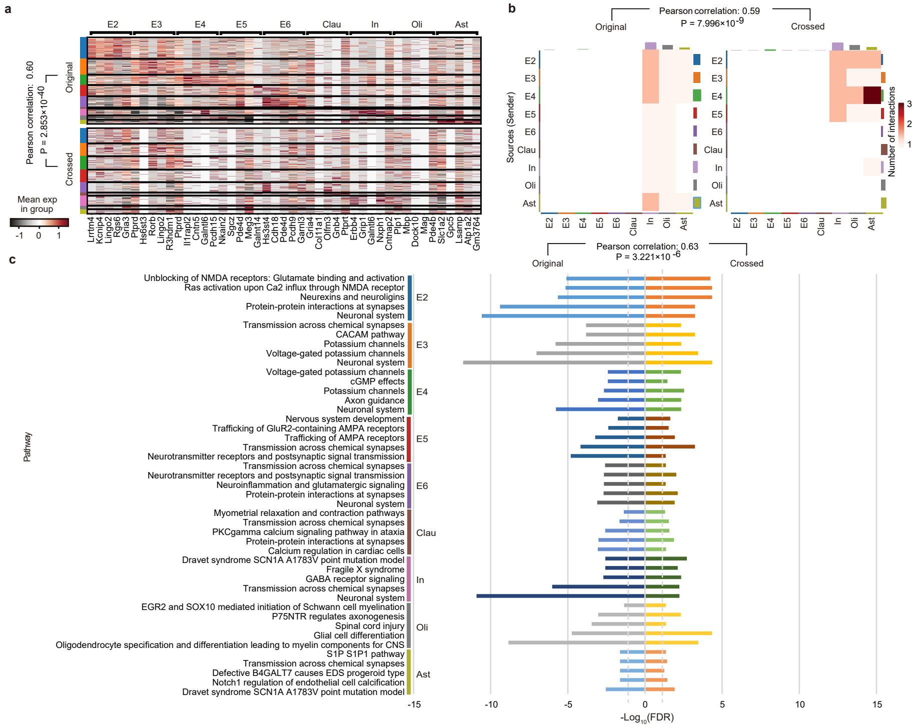

**Supplementary Fig. S6 Cross generation results when the model was trained on independent reference multi-omics datasets. a,** Correlation in Top 5 Gene Expression: The expression levels of the top 5 genes per cell type exhibit a correlation of 0.60 ($P = 2.853 \times 10^{-40}$). **b,** Consistent Cell-Cell Interaction Metrics: Cell-cell interaction metrics consistently align across cell types, with a correlation of 0.59 ($P = 7.996 \times 10^{-9}$). **c,** Validation of Pathway Gene Ratios: Ratios of genes in the most significant pathways for each cell type validate a high correlation of 0.63 ($P = 3.221 \times 10^{-6}$) between the original and cross-generated data.

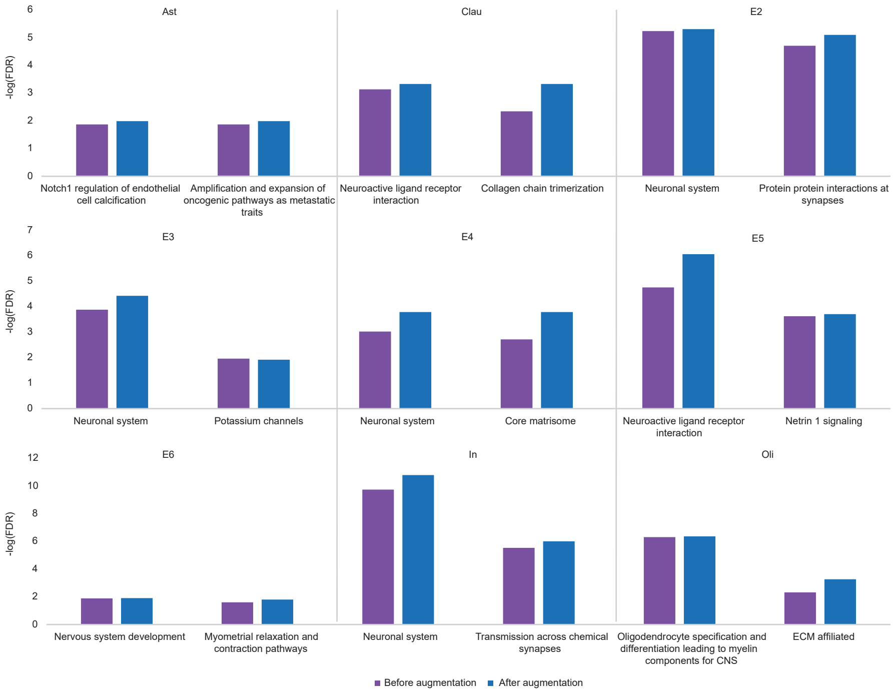


**Supplementary Fig. S7 Augmented scRNA-seq dataset from the matched mouse cortex dataset.** The augmented scRNA-seq data derived from the matched mouse cortex dataset displays an increased capacity to identify a greater number of genes associated with key pathways when compared to the original data. This augmentation is particularly noticeable in each cell type of the matched mouse cortex dataset's scRNA-seq data, which benefits from the integration of scATAC-seq data, allowing for the discovery of more genes related to the key pathways associated with specific cellular functions.

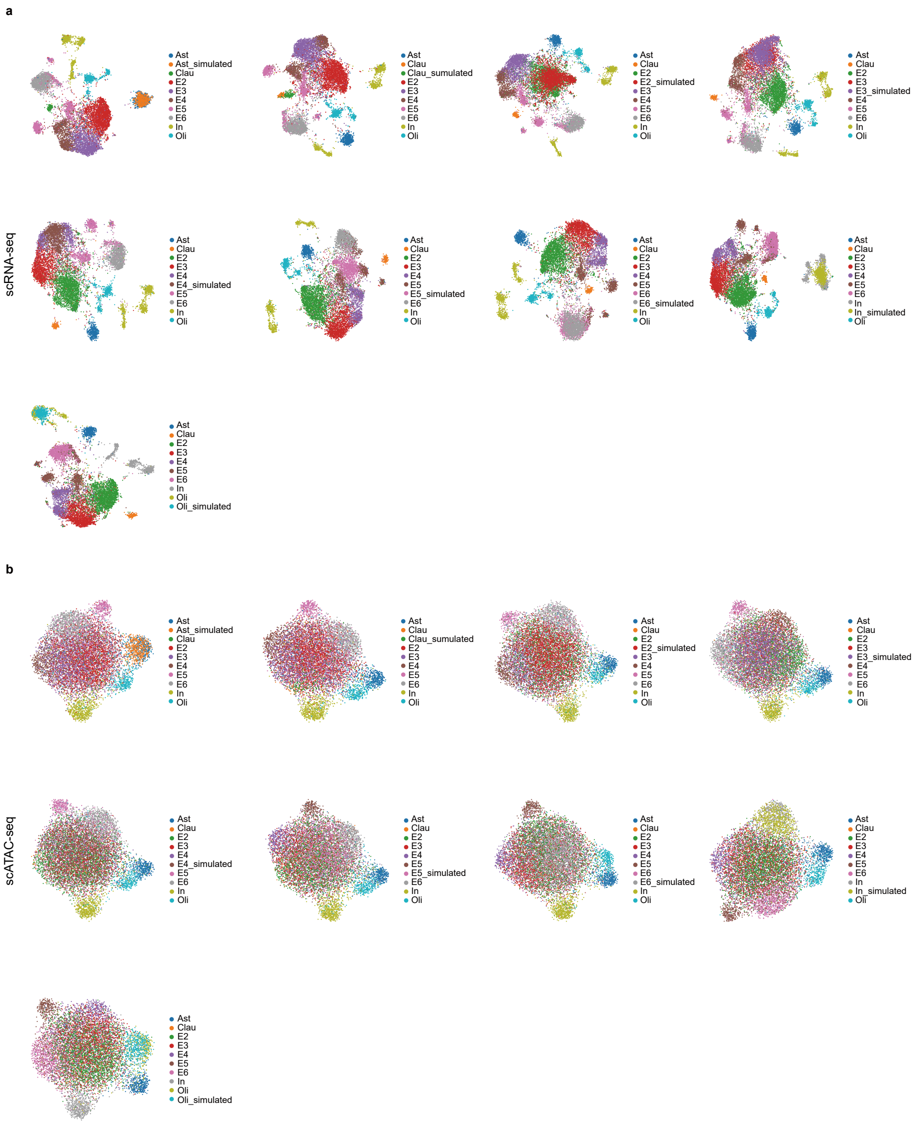


**Supplementary Fig. S8 UMAP of 1X multi-omics simulation of the matched mouse cortex dataset. a,** UMAP of accurate 1X simulation for each cell type in scRNA-seq data of matched mouse cortex dataset. **b,** UMAP of accurate 1X simulation for each cell type in scATAC-seq data of matched mouse cortex dataset.

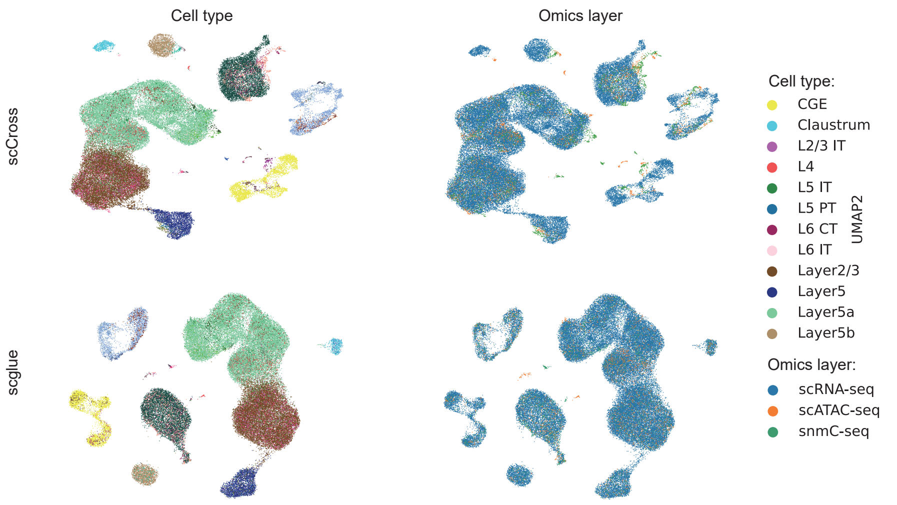


**Supplementary Fig. S9 UMAP visualizations of the cell embeddings in the unmatched mouse cortex dataset compared with different integration methods.** In the UMAP plots, our scCross model clearly outperforms all other integration methods in the unmatched mouse cortex dataset. Besides, we excluded the results of Seurat v4 and uniPort since they do not support triple omics integration.

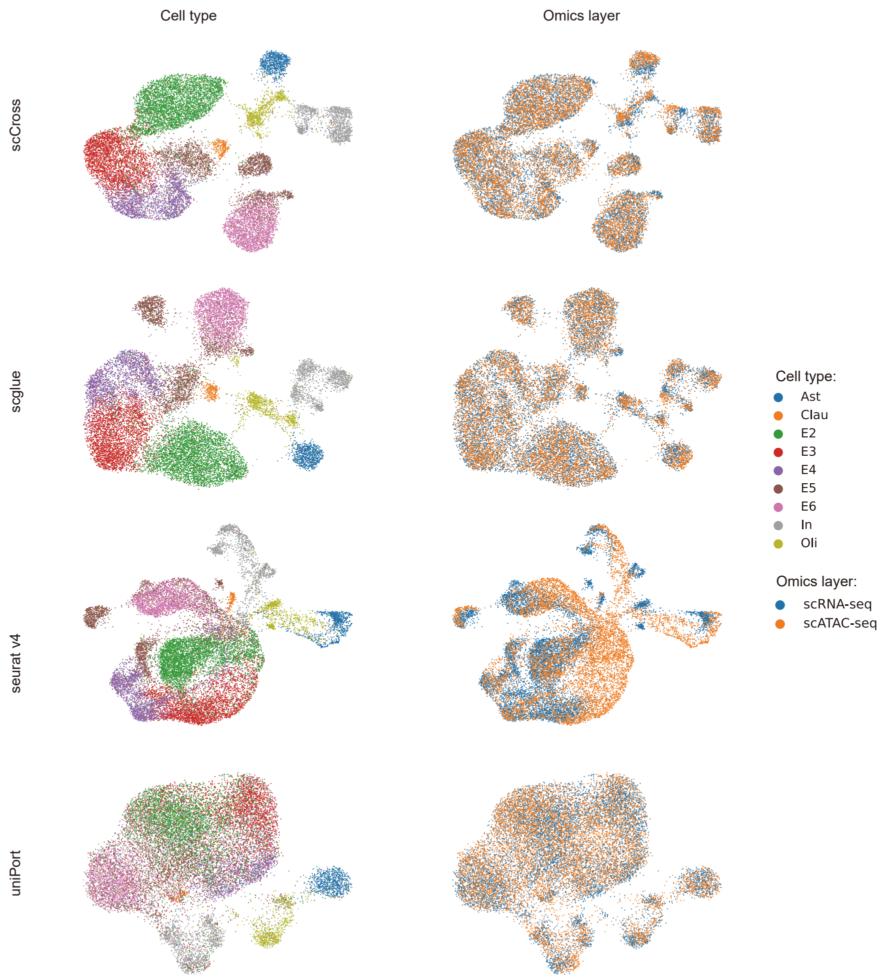


**Supplementary Fig. S10 UMAP visualizations of the cell embeddings in the matched mouse cortex dataset compared with different integration methods.** In the UMAP plots, it's evident that our scCross model outperforms all other integration methods in the matched mouse cortex dataset.

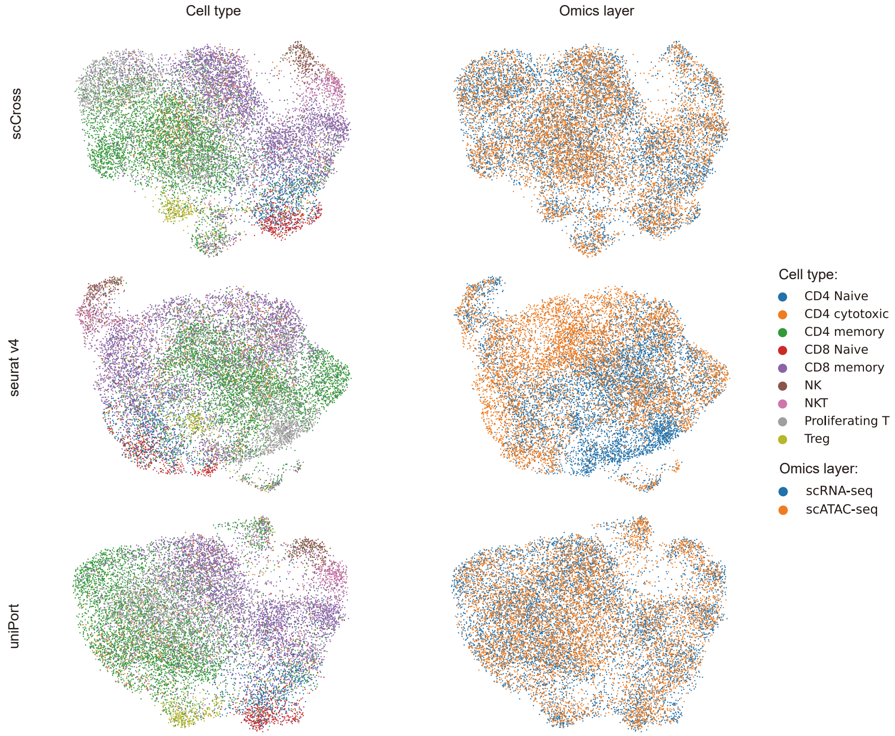


**Supplementary Fig. S11 UMAP visualizations of the cell embeddings in the matched mouse lymphonodus dataset compared with different integration methods.** Evident in the UMAP plots, our model, scCross, surpasses all other integration methods in the matched mouse lymphonodus dataset. Besides, method scglue is unable to produce results in this dataset.

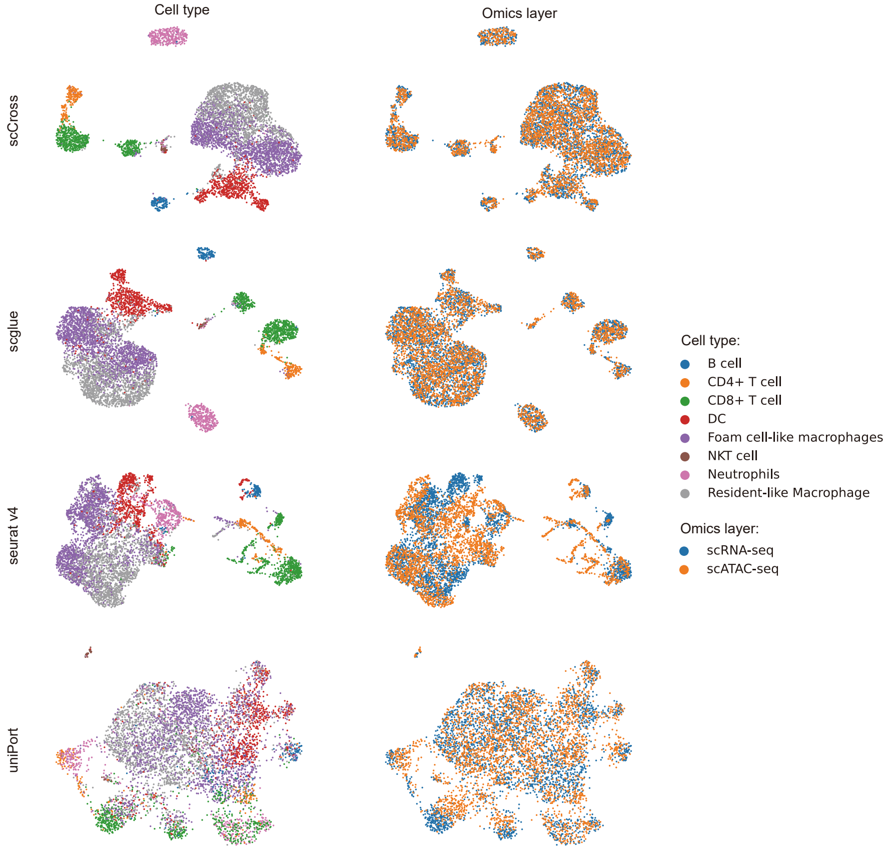


**Supplementary Fig. S12 UMAP visualizations of the cell embeddings in the matched mouse atherosclerotic plaque immune cells dataset compared with different integration methods.** Demonstrated in the UMAP plots, our model, scCross, outperforms all other integration methods in the matched mouse atherosclerotic plaque immune cells dataset.

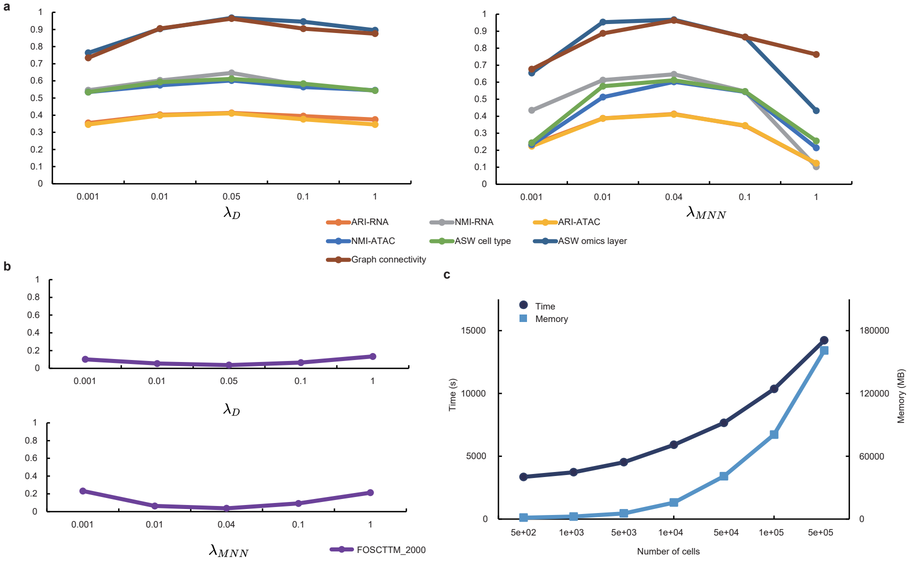


**Supplementary Fig. S13 Time and memory usage, as well as parameter study of scCross. a,** Evaluation metrics, including ARI, NMI, ASW cell type, ASW omics layer, and Graph connectivity performance, were assessed under varying values of $\lambda_D$ and $\lambda_{MNN}$. We performed parameter testing on the matched mouse cortex dataset, and these findings were consistent across other datasets in our study. **b,** The FOSCTTM performance at 2000 cells was analyzed under different values of $\lambda_D$ and $\lambda_{MNN}$. Parameter testing on the matched mouse cortex dataset yielded results that were consistent across other datasets in our study. **c,** We measured the time and memory usage of scCross, observing that our method demonstrates linear resource consumption in both time and memory.

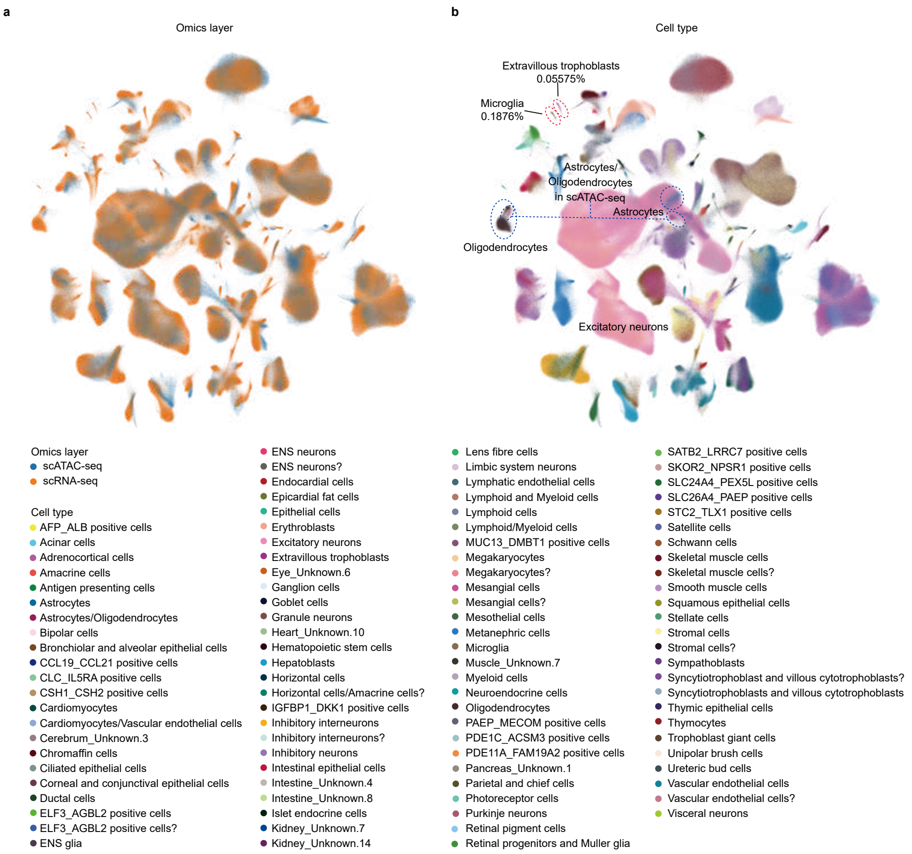


**Supplementary Fig. S14 Integration of the multi-omics human cell atlas by scCross. a,** UMAP visualizations of the integrated cell embeddings, colored by omics layers. **b,** UMAP visualizations of the integrated cell embeddings, colored by cell types. The blue circles pinpoint regions where scCross achieves consistent integration across modalities, resolving discrepancies on the data labels among three omics layers. The red circles spotlight key rare cell populations distinctively identified and integrated by scCross, which were overlooked by scglue.

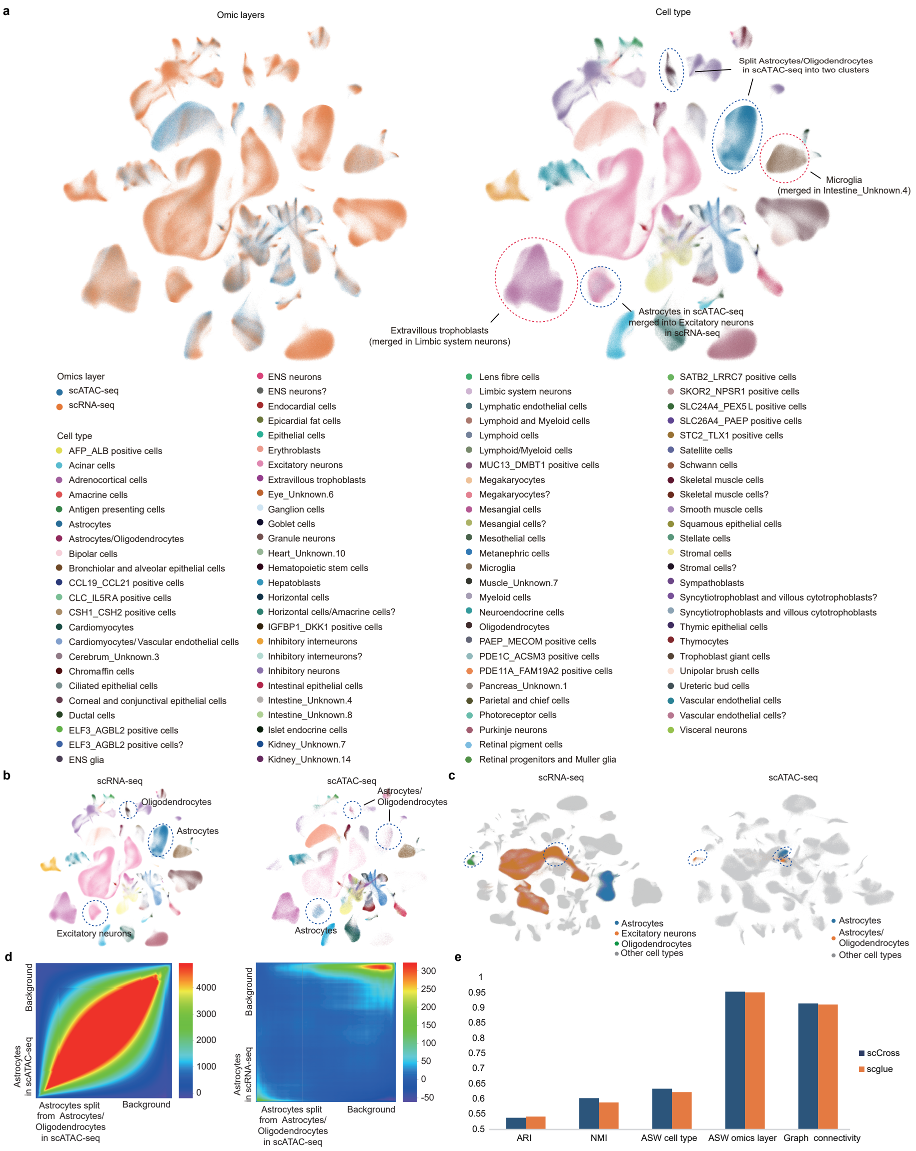
a
Omic layers
Cell type
Split Astrocytes/Oligodendrocytes in scATAC-seq into two clusters
Microglia (merged in Intestine_Unknown.4)
Astrocytes in scATAC-seq merged into Excitatory neurons in scRNA-seq
Extravillous trophoblasts (merged in Limbic system neurons)
Omics layer
scATAC-seq
scRNA-seq
Cell type
AFP_ALB positive cells
Acinar cells
Adrenocortical cells
Amacrine cells
Antigen presenting cells
Astrocytes
Astrocytes/Oligodendrocytes
Bipolar cells
Bronchiolar and alveolar epithelial cells
CCL19_CCL21 positive cells
CLC_IL5RA positive cells
CSH1_CSH2 positive cells
Cardiomyocytes
Cardiomyocytes/ Vascular endothelial cells
Cerebrum_Unknown.3
Chromaffin cells
Ciliated epithelial cells
Corneal and conjunctival epithelial cells
Ductal cells
ELF3_AGBL2 positive cells
ELF3_AGBL2 positive cells?
ENS glia
ENS neurons
ENS neurons?
Endocardial cells
Epicardial fat cells
Epithelial cells
Erythroblasts
Excitatory neurons
Extravillous trophoblasts
Eye_Unknown.6
Ganglion cells
Goblet cells
Granule neurons
Heart_Unknown.10
Hematopoietic stem cells
Hepatoblasts
Horizontal cells
Horizontal cells/Amacrine cells?
IGFBP1_DKK1 positive cells
Inhibitory interneurons
Inhibitory interneurons?
Inhibitory neurons
Intestinal epithelial cells
Intestine_Unknown.4
Intestine_Unknown.8
Islet endocrine cells
Kidney_Unknown.7
Kidney_Unknown.14
Lens fibre cells
Limbic system neurons
Lymphatic endothelial cells
Lymphoid and Myeloid cells
Lymphoid cells
Lymphoid/Myeloid cells
MUC13_DMBT1 positive cells
Megakaryocytes
Megakaryocytes?
Mesangial cells
Mesangial cells?
Mesothelial cells
Metanephric cells
Microglia
Muscle_Unknown.7
Myeloid cells
Neuroendocrine cells
Oligodendrocytes
PAEP_MECOM positive cells
PDE1C_ACSM3 positive cells
PDE11A_FAM19A2 positive cells
Pancreas_Unknown.1
Parietal and chief cells
Photoreceptor cells
Purkinje neurons
Retinal pigment cells
Retinal progenitors and Muller glia
SATB2_LRRC7 positive cells
SKOR2_NPSR1 positive cells
SLC24A4_PEX5L positive cells
SLC26A4_PAEP positive cells
STC2_TLX1 positive cells
Satellite cells
Schwann cells
Skeletal muscle cells
Skeletal muscle cells?
Smooth muscle cells
Squamous epithelial cells
Stellate cells
Stromal cells
Stromal cells?
Sympathoblasts
Syncytiotrophoblast and villous cytotrophoblasts?
Syncytiotrophoblasts and villous cytotrophoblasts
Thymic epithelial cells
Thymocytes
Trophoblast giant cells
Unipolar brush cells
Ureteric bud cells
Vascular endothelial cells
Vascular endothelial cells?
Visceral neurons
b
scRNA-seq
Oligodendrocytes
Astrocytes
Excitatory neurons
scATAC-seq
Astrocytes/ Oligodendrocytes
Astrocytes
c
scRNA-seq
Astrocytes
Excitatory neurons
Oligodendrocytes
Other cell types
scATAC-seq
Astrocytes
Astrocytes/ Oligodendrocytes
Other cell types
d
Background
Astrocytes in scATAC-seq
Astrocytes split from Astrocytes/ Oligodendrocytes in scATAC-seq
Background
4000
3000
2000
1000
0
Background
Astrocytes in scRNA-seq
Astrocytes split from Astrocytes/ Oligodendrocytes in scATAC-seq
Background
300
250
200
150
100
50
0
-50
e
1
0.95
0.9
0.85
0.8
0.75
0.7
0.65
0.6
0.55
0.5
ARI
NMI
ASW cell type
ASW omics layer
Graph connectivity
scCross
scglue

**Fig. S15 Benchmarking on the multi-omics human cell atlas dataset.** **a,** UMAP Visualization of scglue Integration: The UMAP plot reveals that scglue struggles to differentiate small cell clusters effectively (highlighted by red cycles). scglu also find several differences between omics modalities' labels (highlighted by blue cycles), which is detailed in panel b. **b,** UMAP Visualization of scglue Integration in scRNA-seq and scATAC-seq respectively. Here, we can clearly find cells originally annotated as Astrocytes in scATAC-seq are aligned to an Excitatory neurons cluster in scRNA-seq which is verified to be the mislabeling of scATAC-seq data by markers in the Supplementary Fig. S18 of scglue. Beside, the Astrocytes/Oligodendrocytes cluster in scATAC-seq is split into two halves and aligned to the Astrocytes and the Oligodendrocytes clusters (highlighted by blue cycles). **c,** UMAP Visualization of scCross Integration in scRNA-seq and scATAC-seq respectively. scCross can also finds the differences above with scglue. **d,** RROH analysis between the split Astrocytes from the Astrocytes/Oligodendrocytes in scATAC-seq and the Astrocytes in scATAC-seq in left panel and RROH analysis between split Astrocytes from the Astrocytes/Oligodendrocytes in scATAC-seq and the Astrocytes in scRNA-seq in right panel. Here, we use leiden to split the Astrocytes/Oligodendrocytes in scATAC-seq. For fair, we use the common genes between the gene expression and the chromatin accessibility gene activity (2 kb upstream from TSS) in these two analysis. It potentially indicates the inaccuracy of the Astrocytes cell type annotation in scATAC-seq data. **e,** Benchmark Comparison with scglue: In our benchmark analysis of the Human Cell Atlas dataset, our method consistently outperforms scglue. Besides, seurat v4 and uniport do not show their potential in large single cell multi-omics data integration, so we do not benchmark them here.
